# Physical vitrification and nanowarming at human organ scale to enable cryopreservation

**DOI:** 10.1101/2024.11.08.622572

**Authors:** Lakshya Gangwar, Zonghu Han, Cameron Scheithauer, Bat-Erdene Namsrai, Saurin Kantesaria, Rob Goldstein, Michael L. Etheridge, Erik B. Finger, John C. Bischof

**Affiliations:** Department of Mechanical Engineering, University of Minnesota, Minneapolis, USA; Department of Surgery, University of Minnesota, Minneapolis, USA; Department of Biomedical Engineering, University of Minnesota, Minneapolis, USA; AMF Life Systems, LLC, Auburn Hills, USA

**Keywords:** Cryopreservation, Vitrification, Nanowarming, Human Organ, Liter volumes

## Abstract

Organ banking by vitrification could revolutionize transplant medicine. However, vitrification and rewarming have never been demonstrated at the human organ scale. Using modeling and experimentation, we tested the ability to vitrify and rewarm 0.5 – 3 L volumes of three common cryoprotective agent (CPA) solutions: M22, VS55, and 40% EG+0.6M Sucrose. We first demonstrated our ability to avoid ice formation by convectively cooling faster than the critical cooling rates of these CPAs while also maintaining adequate uniformity to avoid cracking. Vitrification success was then verified by visual, thermometry, and x-ray μCT inspection. M22 and EG+sucrose were successfully vitrified in 0.5 L bags, but only M22 was vitrified at 3 L. VS55 did not vitrify at any tested volumes. As additional proof of principle, we successfully vitrified a porcine liver (∼1L) after perfusion loading with 40% EG+0.6M Sucrose. Uniform volumetric rewarming was then achieved in up to 2 L volumes (M22 with ∼5 mgFe/mL iron-oxide nanoparticles) using nanowarming, reaching a rate of ∼88 °C/min with a newly developed 120 kW radiofrequency (RF) coil operating at 35kA/m and 360kHz. This work demonstrates that human organ scale vitrification and rewarming is physically achievable, thereby contributing to technology that enables human organ banking.

## Introduction

Vitrification, or the rapid cooling of biologic material to an ice-free glassy state at ultralow temperature, promises indefinite storage of cells, tissues, and organs for transplantation or other biomedical applications [1, 2]. Over a century ago, the first successful vitrification of frog spermatozoa was conducted by Luyet and Hodapp (in 1938) [3, 4]. Subsequently, by the 1980s, vitrification was performed on multiple biological systems, including whole rabbit kidneys (10’s of mL volumes) [1]. Since then, vitrification has been attempted up to 1.5L but failed due to underlying ice formation and fracture [5] inherent to heat transfer at these scales. Even under conditions where vitrification upon cooling was successful, even faster rates are required upon rewarming, leading to failures (ice formation and/or cracking) at the rabbit kidney scale [6]. The critical cooling rate (CCR) and critical warming rate (CWR) are the required cooling and warming rates needed for vitrification and rewarming without ice formation.

Several volumetric rewarming techniques have since been tested to overcome the limitations of conventional boundary layer rewarming, including high-intensity focused ultrasound (HIFU) and dielectric/microwave warming. HIFU heats by focusing sound/pressure waves on the vitrified material but is currently limited to volumes ≤ 2 mL [7]. It also has the inherent limitation of wave penetration, reflections, acoustic impedance mismatch at interfaces, cavitation, and the possibility of thermal overshoot in the liquid phase [8, 9]. Dielectric/microwave rewarming (heating of dipolar molecules by electromagnetic (EM) waves) potential rewarming volumes can approach ∼100 mL in CPA systems [10], including rabbit kidneys (∼50 mL) [11]. However, this approach also has several limitations, including small penetration depths, poor EM coupling in the vitrified “solid” state, and the potential for “thermal runaway” near the melt [12], each of which are accentuated by the inhomogeneity typical of biological tissues (and their associated dielectric properties), all of which are expected to worsen as the scale increases.

To address these limitations, our group developed a volumetric rewarming technology termed “nanowarming,” where magnetic iron-oxide nanoparticles (IONPs) are perfused through the organ vasculature before vitrification. For rewarming, the organs are placed in an alternating magnetic field that couples to the nanoparticles and induces heating from within the vasculature through magnetic hysteresis. Since the chosen radiofrequency waves penetrate the system without significant attenuation, this approach is, in theory, fully scalable to human organs. So far, we and others have published on nanowarming of volumes up to 80 mL CPA mL and 30 mL biological nanowarming at rates up to 50 - 100 °C/min [13-18] (see Table S1).

To achieve cryopreservation of human organs through vitrification, we anticipate needing 0.5 - 1 Liter volumes for hearts and kidneys, while human livers may require ≥ 3 L [19]. However, neither successful vitrification nor rewarming has been demonstrated in the literature at these scales. Since capillary spacing is relatively conserved across organs and tissues, uniformity and heating rate depend primarily on the concentration of IONPs, distribution of the magnetic field strength, and frequency. Given the timescales of rewarming, heating is practically uniform if the magnetic field is uniform as the heat rapidly diffuses across intercapillary distances.

There are currently no volumetric means of cooling human organs, so convective cooling will be directly impacted by the size of the system, with the cooling rate at the center of the sample decreasing as the size of the system increases [20]. This may lead to insufficient cooling rates and/or physical fractures/cracking due to thermal gradients in the vitrified state [21] if a proper cooing protocol is not employed. It is also important to note that since nanowarming rates are independent of sample volume, cooling conditions will become the rate-limiting step in vitrifying and rewarming human organ scale volumes.

Here, we describe the conditions for successfully achieving physical vitrification and nanowarming in human organ scale volumes. Fig. 1 shows the sequential steps for this multi-liter vitrification and CPA nanowarming in a cryobag. We report the successful vitrification of volumes up to 3L and uniform nanowarming from the vitrified cryogenic state of up to 2L. We further demonstrate physical vitrification in a CPA-perfused liver (∼1L porcine liver). This study demonstrates that bulk systems equivalent in size to human organs can be vitrified and nanowarmed, as verified by visual inspection, thermometry, and μCT. Combined with our group’s related demonstration of a robust protocol for long-term cryopreservation, rewarming, and successful transplantation of a functional preserved rat kidney [14], these findings lay the groundwork for achieving clinical organ banking.

**Fig. 1:**
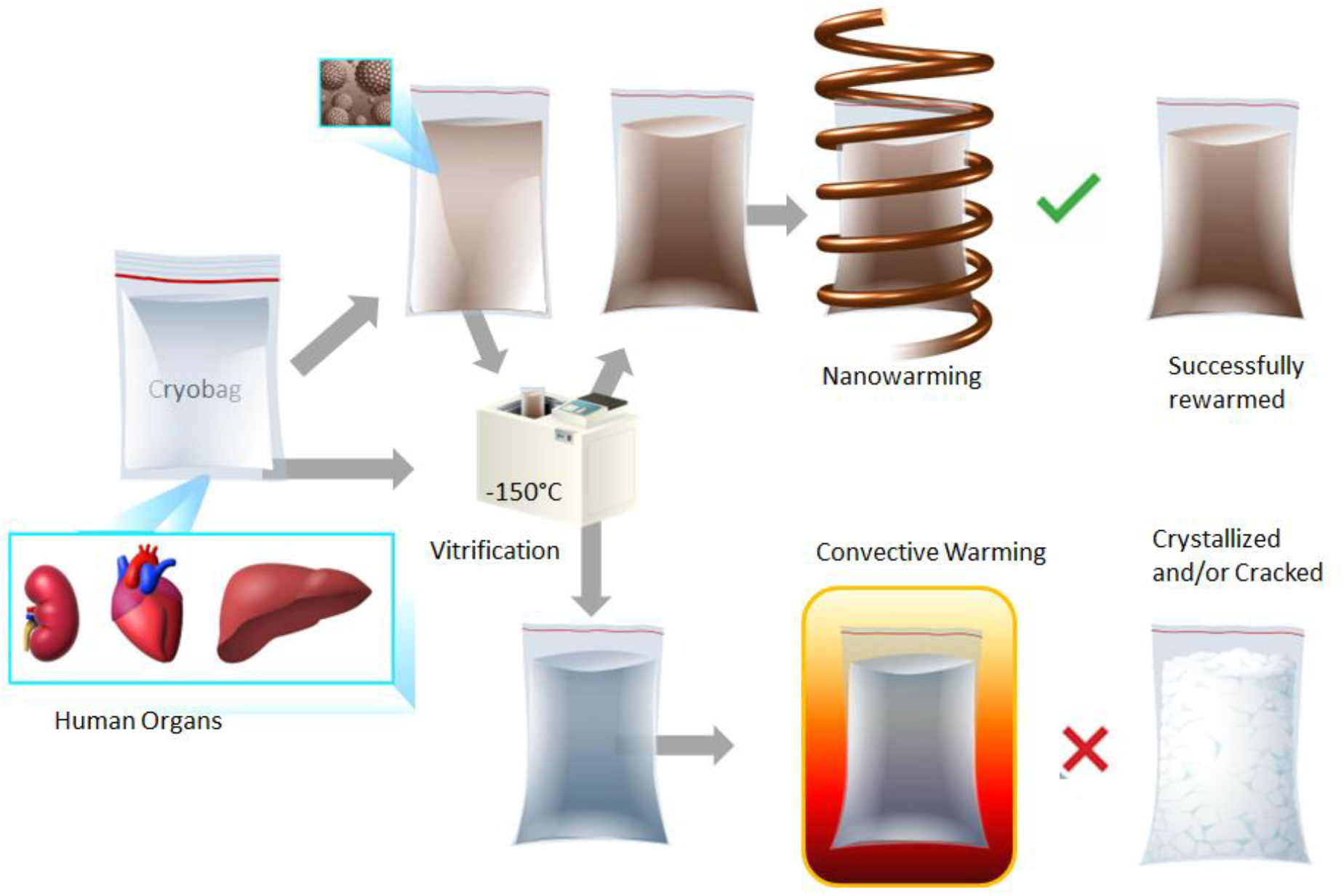
Schematic flow of steps (left to right) in liter scale vitrification and rewarming. Liter volumes of a CPA (0.5-3L) in cryobags are large enough to hold a human organ. The cryobag is placed inside a controlled rate freezer (CRF) for cooling. For nanowarming (top section of the flow chart), the cryobag is vitrified with iron oxide nanoparticles (IONPs) suspended in the CPA. The vitrified cryobag is placed inside the RF coil, alternating magnetic fields are turned on, and the IONPs generate heat. This leads to successful (rapid, uniform) rewarming, avoiding crystallization or cracking failure modes even at the liter scale. Traditional rewarming employs convection using a water bath, which at these scales results in ice recrystallization and/or fractures due to insufficient rewarming rates (slow) and thermal stresses (non-uniform), respectively (lower section of the flow chart). IONP, iron oxide nanoparticles; CPA, cryoprotective agent; CRF, controlled rate freezer.

## Results

### Modeling Liter Scale Vitrification

Computational heat transfer modeling was performed to develop the vitrification protocols of 0.5, 1, and 3 L cryobag systems using COMSOL 5.4. Convection within the CPA itself is neglected due to the high viscosity of CPAs at subzero temperatures. Therefore, the general form of the governing equation for cooling and warming is assumed to be the conduction heat equation:

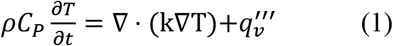

Where ρ is density, C_p_ is specific heat, k is thermal conductivity, T is temperature, and q_v_’’’ is volumetric heating (generated from IONP = volumetric specific absorption rate (SAR_V_)). Limited data on CPA thermal properties are available, hence we assume them to be that of M22 (listed in Table S2) [22], which should be a reasonable approximation for each of the other CPAs. The initial CRF temperature was set to 0 °C (see Fig. S4). Boundary cooling at h = 100W/m^2^K (h: heat transfer coefficient) and T_CRF_ ∼-40°C/min (maximum CRF cooling rate) was initiated, and upon reaching a temperature of -122°C, just above T_g_, the system was annealed (thermally equilibrated to minimize thermal gradients/stress, Fig. S4). Two important modeling outcomes studied were the cooling rate (dT/dt) in the region of ice formation (above ∼-100°C), and the temperature difference (ΔT= T_center_-T_edge_) in the glassy region (below T_g_: glass transition temperature) where the system is at most risk of fracture/cracking failure. The characteristic length for heat transfer is defined as L_C_ (=Volume/Surface Area) [23], which can be used to compare the trends in thermal predictions across different volumes (see Fig. S4). Note that larger volumes (and L_C_) result in slower cooling rates and require longer annealing time (Table 1). After annealing, the system was cooled slowly (<1 °C/min) to -150°C for storage to minimize thermal stress, keeping the temperature difference (< 20 °C) in the brittle glassy phase [20, 24]. The summary of vitrification success for all 3 volumes and CPAs is shown in Fig. 2 (and Fig. S10). The temperature distribution across the geometry is plotted for the three CPA volumes (Figs. 3, S3 and S12). The minimum predicted and measured cooling rate (CR) (which is at the center) decreased with increased volumes from ∼1.4°C/min for 0.5L (L_C_∼1.2cm) to ∼1°C/min for a 1L (L_C_∼1.4cm) and finally ∼0.5°C/min for 3L (L_C_∼2.2cm) cryobag.

**Table 1:** Summary of cooling protocols developed based on heat transfer modeling for each vitrification volume.

| CRF Cooling Protocol | 0.5 L | 1 L | 3 L |
| --- | --- | --- | --- |
| Characteristics Thermal Length: $L_c$ | 1.2 cm | 1.4 cm | 2.2 cm |
| Start Temperature | 0°C | 0°C | 0°C |
| Chamber initial cooling rate (to -122°C) | -40°C/min | -40°C/min | -40°C/min |
| Anneal (hold) at -122°C | 180 min | 250 min | 520 min |
| Chamber cooling rate in glassy region (from anneal to -150°C) | -0.5°C/min | -0.4°C/min | -0.2°C/min |
| Hold at -150°C | 35 min | 50 min | 90 min |
| Total Protocol Time (hrs) | 4.6 | 6.2 | 12.5 |
| Modeled Center Cooling Rate (0 to -100°C) | 1.36 | 0.96 | 0.45 |

**Fig. 2:**
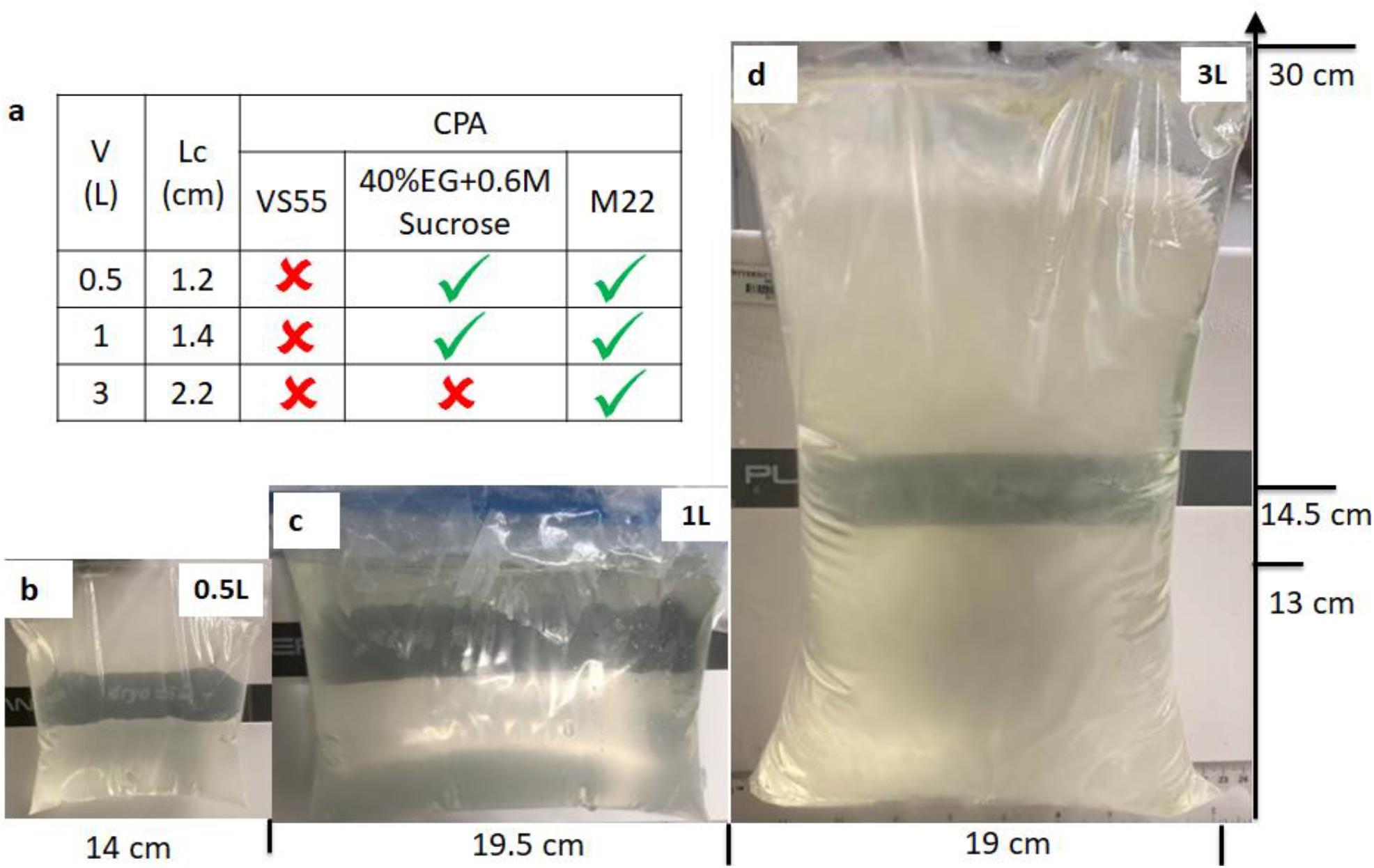
Demonstration of physical success of vitrification in multiple volumes. **a** Table summarizes vitrification results for all the 3 CPAs and volumes. Photos of a successful vitrified (glass) M22 inside a cryobag for **b** 0.5 Liter, **c** 1 Liter, and **d** 3 Liter (largest volume reported). The out-of-plane thicknesses are 5.5, 6.5, and 10.5 cm for 0.5, 1, and 3L cryobags, respectively.

**Fig. 3:**
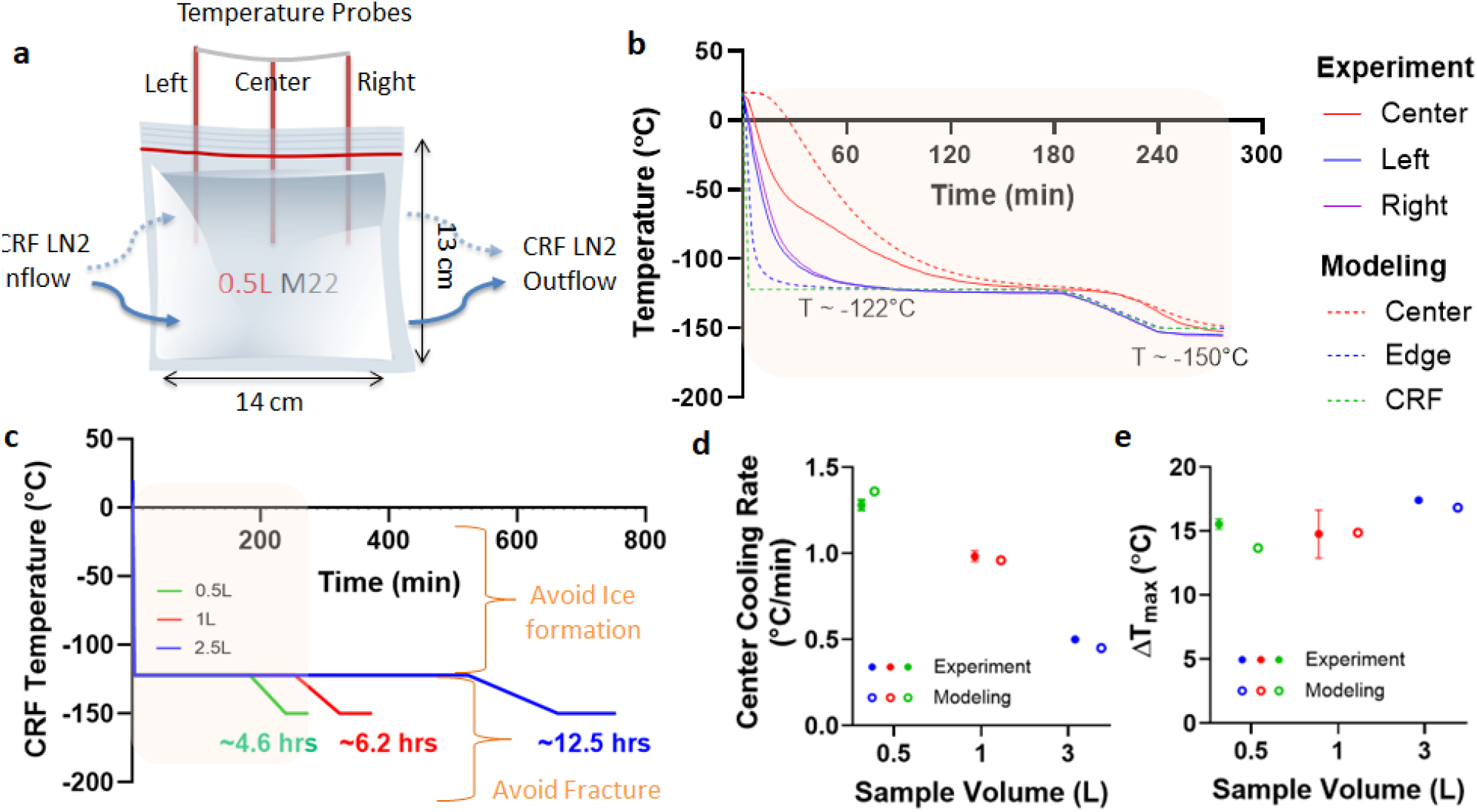
Thermal results from experimental and modeled liter scale CPA vitrification. **a** Schematic for a representative case, 0.5 L cryobag containing CPA with placement of three fiber optic temperature probes (3 cm apart). Blue arrows show the direction of LN2 flow in CRF. **b** Experimental and predicted temperature vs. time plot for 0.5L M22. The dashed green line shows the programmed CRF temperature profile/protocol. **c** CRF cooling protocols for 0.5, 1, and 3 L volumes. The regions of ice formation and fracture danger are labeled. Scatter plot of **d** center cooling rate and **e** temperature difference (ΔT_max_ in the glassy region) for all three volumes tested for M22 (mean ± SD; n=3). Cooling rate is calculated in ranges 0 to -100 °C and -120 to -150 °C for temperature difference plots. Mean cooling rates are greater than the CCR of M22 (∼0.1°C/min). Temperature differences are within the allowable limit (dashed) (< 20°C) calculated from a simple thermal shock equation [20].

### Measuring Liter Scale Vitrification

Based on these cooling protocols (Fig. 3C), we then cooled cryobags containing three different CPAs (M22, VS55, 40%EG+0.6Msucrose) at the three modeled volumes (0.5, 1, and 3L). The success or failure of vitrification in these systems was verified by visual, thermal, and μCT measurements. Visually, ice could be identified as round spherulites during crystallization (Fig. S8A) or by opaque, milky-white/cloudy appearance throughout the sample in case of complete crystallization (center of Fig. S8B, S8C). Another mode of physical failure is fractures or cracking, which could be visually observed as the presence of linear defects (Fig. S9). In the absence of failure modes (crystallization and fractures), the CPAs looked clear, transparent, and glassy, implying successful vitrification (Fig. 2). In the case of organs, failure was assessed visually on the surface and internally by photos of the bisected vitrified organ to evaluate for ice. Vitrified material is more radiodense (higher Hounsfield Units, HU) on μCT than ice, and cracks could be detected directly by abrupt changes in radiodensity [13, 16] (Fig. S13).

Successful vitrification in M22 was achieved for all volumes tested (Fig. 2). 40%EG+0.6Msucrose was also successfully vitrified with no visual ice formation up to 1L (Fig. S10). Ice formation was observed in VS55 for all three volumes (Fig. S8 and S10). This was expected as the achieved cooling rate for these three volumes (0.5L∼ 1.4°C/min, 1L∼1°C/min, 3L∼0.5°C/min) were lower than the CCR of VS55 (∼2.5°C/min). We further confirmed our visual findings using μCT, which showed successful vitrification in M22 and 40%EG+0.6Msucrose but ice formation in VS55 at 0.5L volumes (Fig. S13). The temperature profiles, cooling rates, and temperature differences are shown in Fig. 3 (and Fig. S12). As before, the cooling rates decreased, and the temperature differences increased with increasing sample volume.

### Vitrification of a human-scale organ - porcine liver

To test our ability to vitrify an organ at these scales, we perfused a ∼800 mL porcine liver (similar in size to juvenile, age ∼ 10-year-old, liver and greater volume than most other adult human organs) with 40%EG+0.6Msucrose. We cooled it in a cryobag with a surrounding CPA volume of ∼ 200 mL. The 0.5L CRF cooling protocol (Fig. 3C) was used for the liver because the estimated L_C_ (∼0.9 cm) of the porcine liver in CRF was closer to the L_C_ (∼1 cm) of the representative 0.5 L cryobag volume (see Fig. S7). This highlights the importance of appropriately designing cooling protocols for a given geometry. The resulting vitrified porcine liver is shown in Fig. 4. The liver appeared vitrified based on visual inspection, except for a small amount of ice formed around the portal vein and fatty tissue (poorly vascularized perihilar tissue), which presumably did not equilibrate fully with the CPA. The vitrified liver was also cross-sectioned (Fig. S14) and largely appeared to be free of ice. Modeling predicted the center cooling rate of the liver as ∼4°C/min, which exceeds the CCR (∼1 °C/min) of 40%EG+0.6Msucrose, which further supports vitrification (see Fig. S6).

**Fig. 4:**
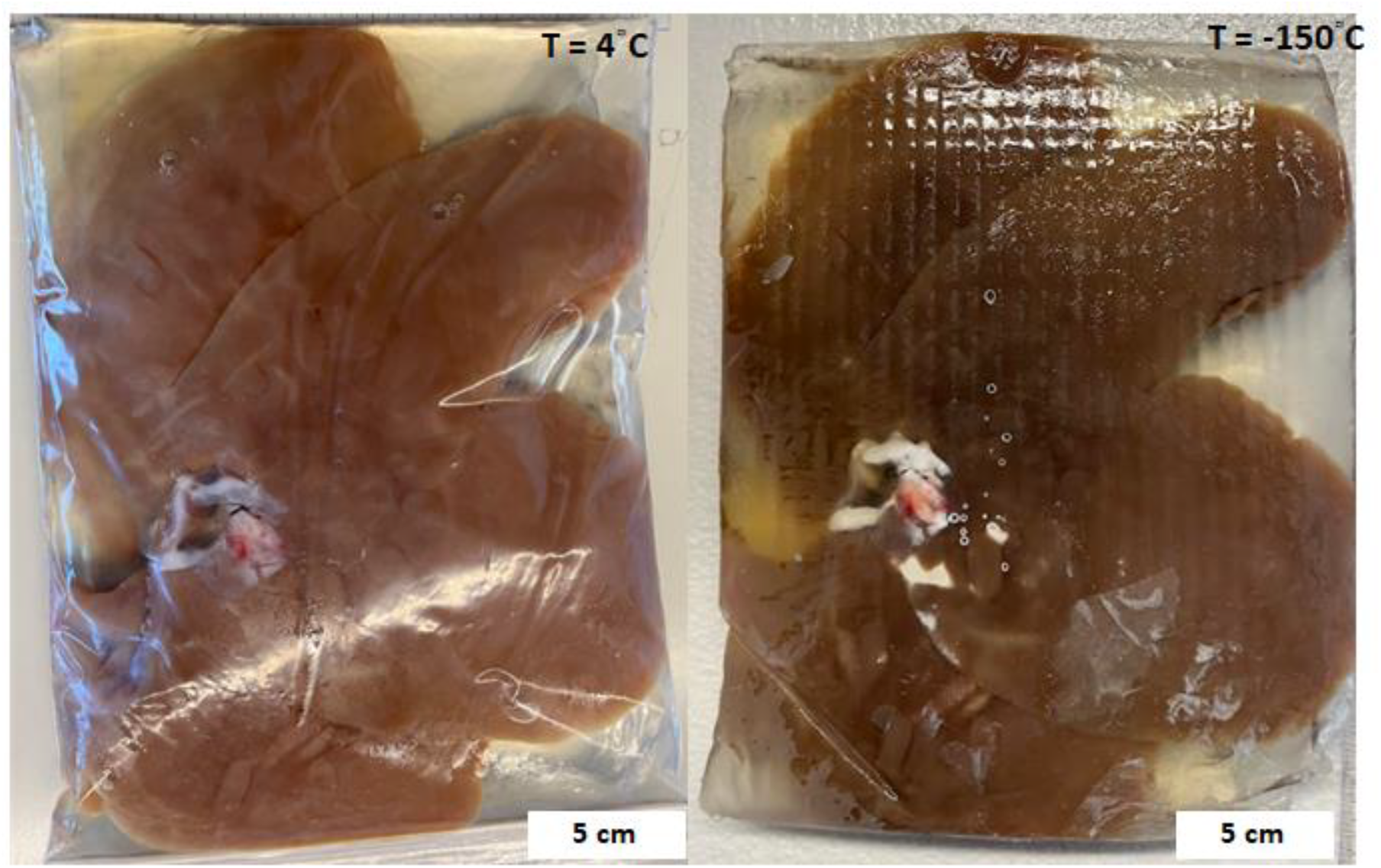
Photos of the porcine liver (left) before (T = 4°C) and (right) after vitrification (T = -150°C). The pattern in the photo was due to the cryobag placement on a supporting mesh in the control rate freezer (CRF) (see Fig. S7B). The cryobag was removed for the vitrified liver photo to reduce glare and get a clear photo.

Effective cryobag sealing was beneficial in preventing open-surface ice formation and reducing the size of the system (L_C_). Vacuum sealing with the removal of surrounding CPA solution led to ice on superficial surfaces of the liver, potentially due to cryobag surface nucleation, so having a layer of CPA around the liver in the bag proved beneficial for successful vitrification.

### Characterization of liter scale RF rewarming coil

We next evaluated the ability to uniformly rewarm liter-scale volumes from the cryogenic vitrified state (< -120°C). Although past studies have rewarmed whole rat and rabbit organs using a 15 kW RF coil (AMF Life Systems) [13, 16, 18] or other commercial RF coils [15, 17], these studies were typically limited to < 30 mL, with only a few nearing ∼100 mL volumes (Table S1) [6]. As noted earlier, uniformity in rewarming will depend on the uniformity of the applied field, assuming practically uniform IONP distributions (i.e., through vascular perfusion to the capillary beds). Therefore, to achieve uniform nanowarming in clinical scale volumes, we worked with AMF LifeSystems, LLC. to design, build, and characterize a 120 kW RF coil with a 2.5L uniform field region. This substantially extends our capabilities beyond previous nanowarming studies (Fig. 5).

**Fig. 5:**
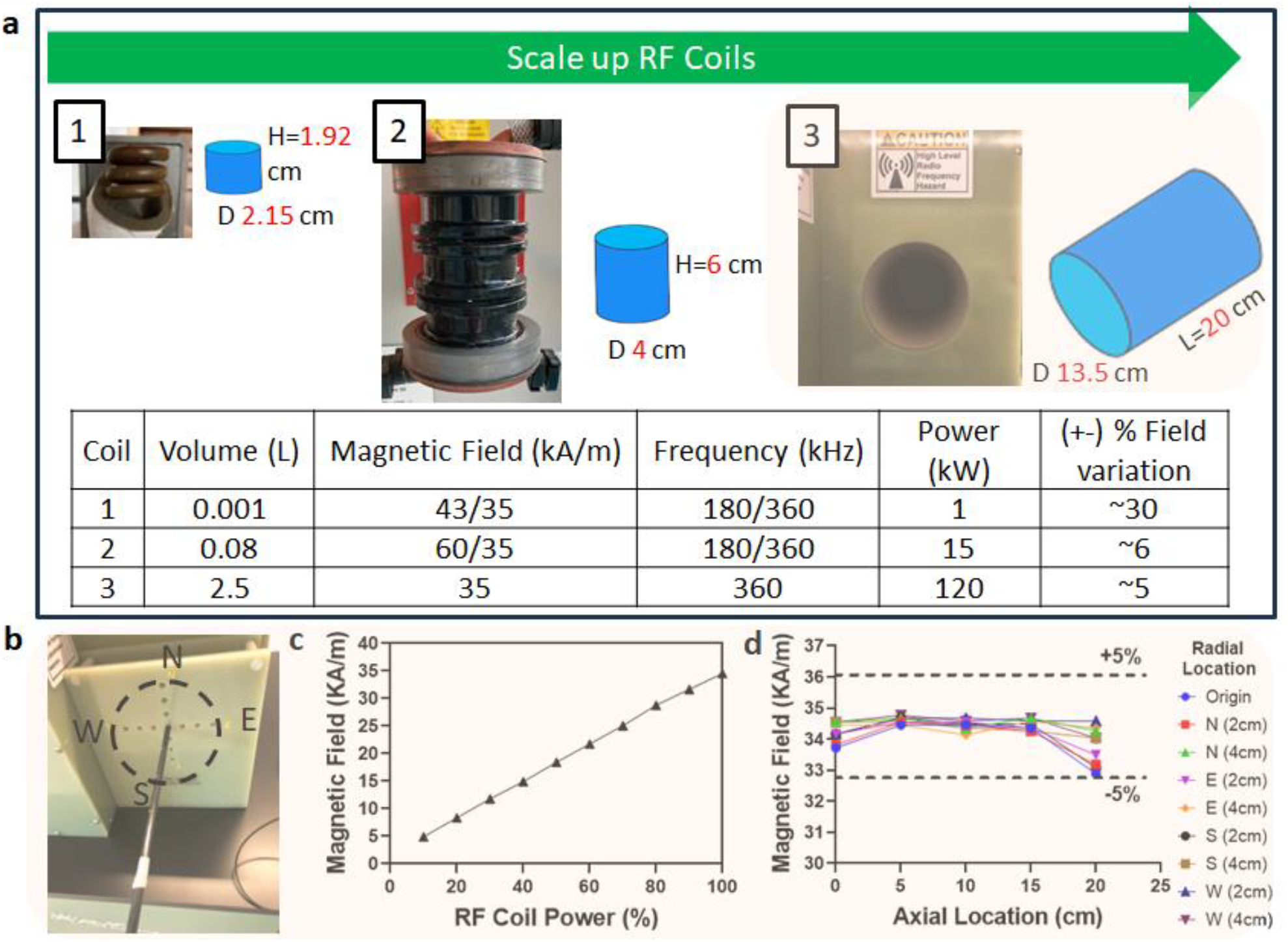
Comparison of several generations of radiofrequency (RF) coil systems and characterization of 120kW RF coil for liter scale nanowarming. **a** Photo of the RF coil system is shown alongside the schematic of the coil’s approximate uniform volume (blue cylindrical region) with diameter (D) and length (L) listed next to blue region (not to scale). The bottom table compares operating power, magnetic field strength, frequency, sample volume, and uniformity of the systems. Note that the RF coil uniformity is a % variation of magnetic field strength across the RF coil volume (blue cylinders). The top green arrow shows the increasing sample scale for RF coils with practical capacities, such as 1mL cryovials in a 1kW coil, human ovaries and eyes in a 15kW coil, and human kidney and heart in a 120kW coil. **b** Photo of the RF coil bore and RF probe placement for magnetic field measurements with labels showing relative radial directions (North (N), South (S), East (E), and West (W)). **c** Plot of measured magnetic field strength vs. coil power. **d** Plot of magnetic field strength vs. axial location for different radial locations. The 120 kW system frequency is fixed at 360 +/-5 kHz.

Detailed characterization of the alternating magnetic field within the RF coil was performed across the volume of interest in the new 120 kW system (Figs. S15-17) and compared to previous coil systems [6, 25]. We measured the magnetic field using a 2D RF probe coil (AMF Lifesystems, LLC.) as a function of spatial distribution and applied coil voltage. At full power, the RF coil generated up to 35 kA/m magnetic field strength across the coil’s volume at a frequency of 360 kHz. The calibration showed a linear relationship between applied power and field strength (Fig. 5C) for the 120 kW coil. The magnetic field was spatially uniform (within 5%) for a 20 cm axial distance and 6cm radial direction (Fig. 5D). Importantly, the maximum field strength and frequency output were comparable between the 15 kW and 120 kW systems. Additional power in the larger RF coil was required to sustain this field uniformly across the substantially higher coil volume (∼80 mL versus 2.5 L).

To best design nanowarming protocols for use in the new system, we systematically evaluated IONP heating across the range of available field strengths, frequencies, and temperatures. To accomplish this, we measured the specific absorption rate (SAR_Fe_, normalized to IONP mass) for IONPs in CPA (M22) as a function of magnetic field and frequency at both room and cryogenic temperatures. As seen in Fig. 6A, for the IONPs tested, SAR increased with magnetic field strength, though it saturated at higher field strengths, as observed elsewhere [26]. SAR_fe_ increased at higher frequencies (∼2 times more at 360 kHz than 190 kHz for 35 kA/m), suggesting a proportional increase in heating with frequency, consistent with prior reports [25, 27, 28]. Given that the output field is capped by the power available to the system (P ∝ freq. x H), proportionally higher rewarming is expected at higher frequency and field strength (e.g., 360 kHz and 35 kA/m vs. 180 kHz and 60 kA/m). Lastly, we report that SAR_V_ increased at cryogenic temperature (by ∼ 1.5 times) compared to room temperature (Fig. 6B). Further details of the SAR calculation can be found in the supplementary information (also see Fig. S18 for SAR of uncoated IONPs: EMG308).

**Fig. 6:**
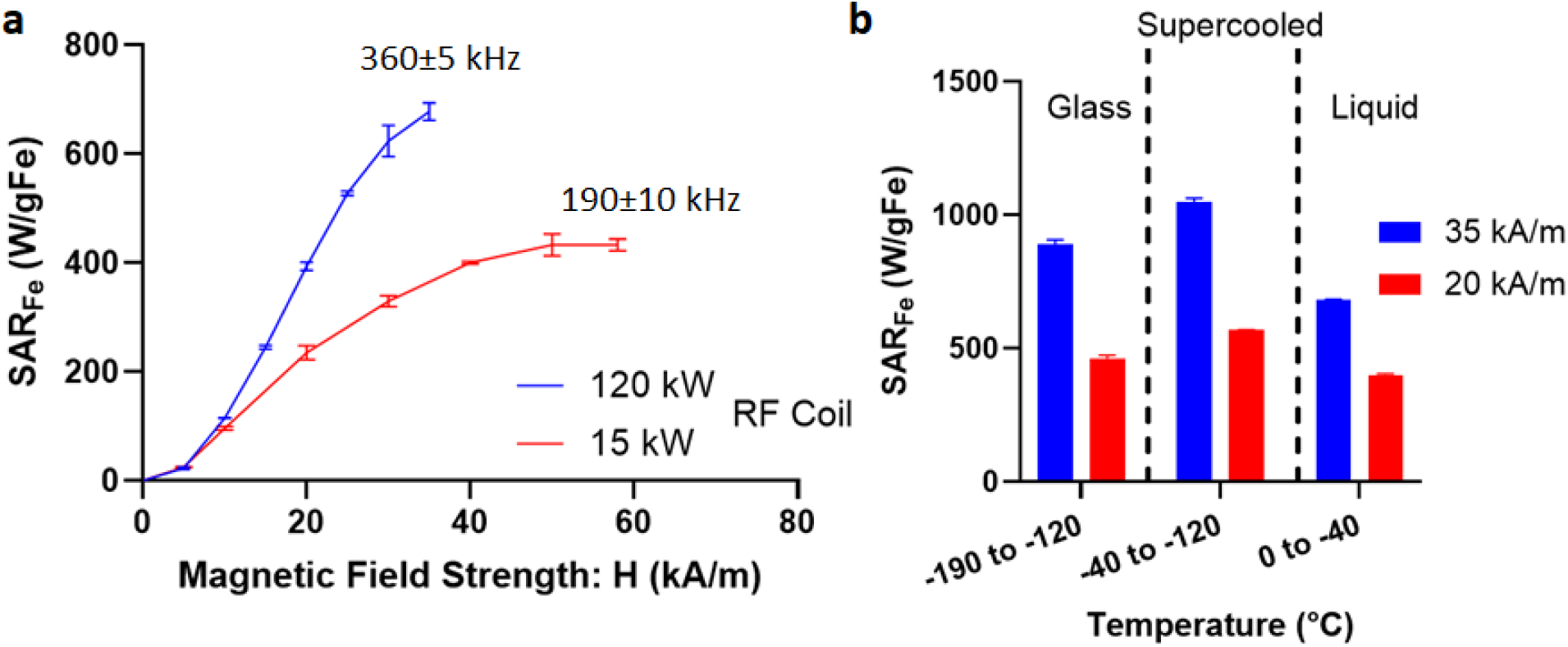
Nanowarming specific absorption rate (SAR). **a** Plot of SAR_Fe_ (SAR_V_/ C_Fe_) vs. magnetic field strength (H) measured at room temperature for iron-oxide nanoparticles IONPs (sIONPs in M22 shown here) at two frequencies (190 and 360kHz) (plotted mean ± SD; n=3). **b** Plot of SAR_Fe_ vs. temperature for sIONPs in M22. Average SAR_Fe_ (mean ± SD; n=3) is plotted in three different temperature regions, i.e., glass, supercooled, and liquid. SAR is measured from cryogenic temperature (−196°C) to room temperature (20°C) at two different field strengths (20, 35 kA/m) and 360 kHz (see Fig. S18 for more details).

### Liter scale nanowarming

To demonstrate nanowarming scales relevant for vitrified human organs, we prepared M22 with IONPs at ∼10.7mgFe/mL for 1L and ∼4.6 mgFe/mL for 2L volume. Different cryobags than those used in the vitrification studies were used to fit in with the workable volume of our 120 kW RF coil (Fig. 7A, also see Fig. S20). After the 1L and 2L volumes were vitrified in the CRF following the same protocols described above, they were stored in a -150°C freezer overnight. For rewarming, the samples were rapidly transferred to (within ∼3-5 seconds) and rewarmed inside the 120 kW RF coil. Samples were rewarmed to 0°C within ∼1min for 1L and ∼2min for 2L at rates of ∼172°C/min and ∼88°C/min, respectively (Fig. 7C). This difference is due to the use of ∼50% lower IONP concentration in the larger 2 L bag. Rates were identical when 0.5L and 1L volumes were rewarming at the same IONP concentration (Fig. S21). In all cases, the measured temperature differences across the cryobag volume were negligible (∼< 5°C) (Figure 7D).

**Fig. 7:**
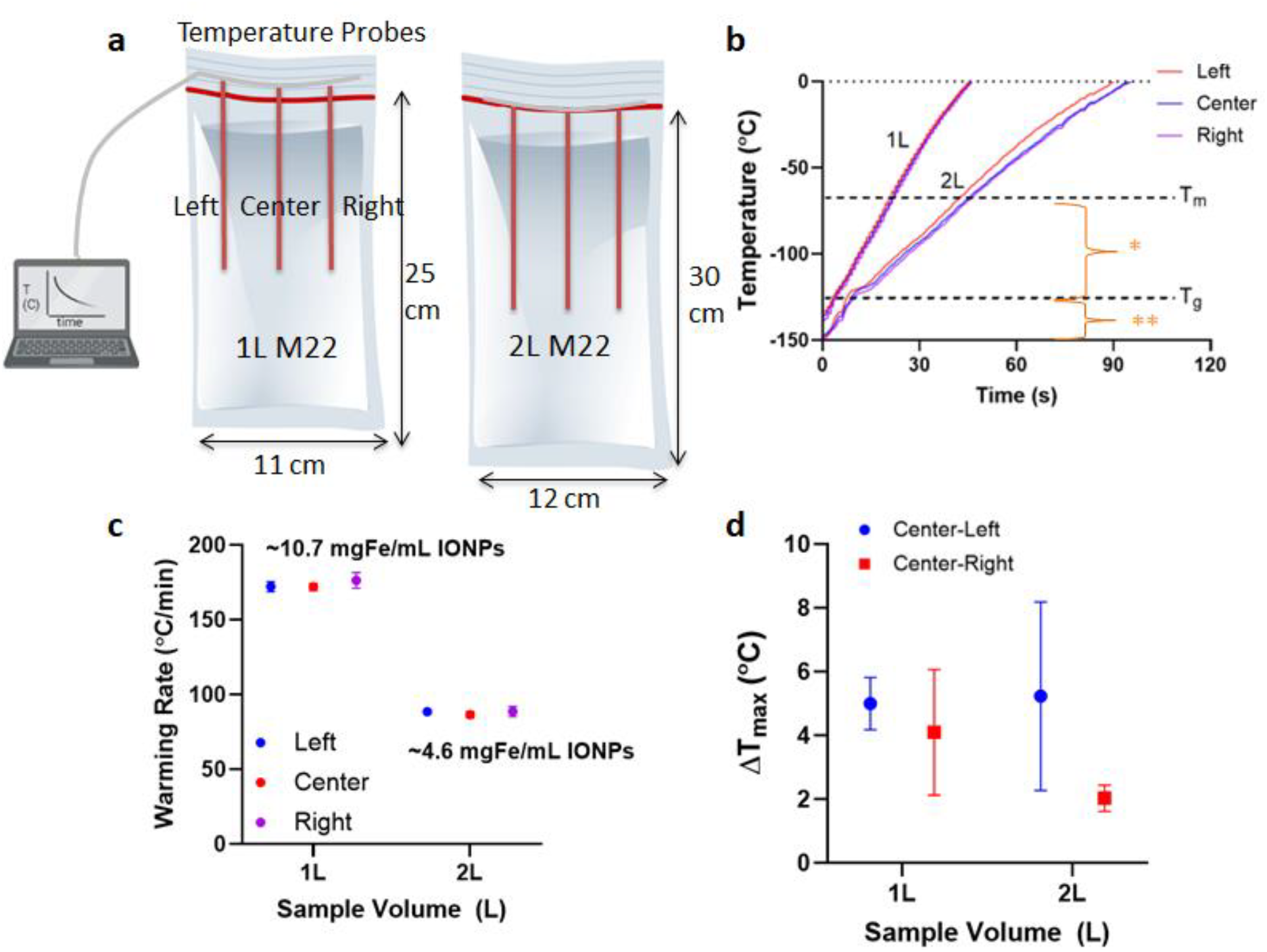
Physical demonstration of liter scale nanowarming. **a** Schematic showing 1L (left) and 2L (right) cryobag containing CPA with three fiber optic temperature probes (placed 4 cm apart / 13 cm depth inside 1 L and 15 cm depth inside 2 L cryobags). The dimensions of the cryobag are shown. **b** Temperature vs. time plot for rewarming of 1L and 2L volumes of M22 (differences in rates due to relative differences in IONP concentrations). Region with greatest risk of ice formation is shown as (*) and fracture failure as (**). Scatter plots of **c** average rewarming rate and **d** temperature difference (∆T) for 1 L and 2 L M22 (mean ± SD; n=3). The rewarming rate is calculated from 0 to -100°C. Average rewarming rates are greater than the CWR of M22 (∼0.4°C/min). Maximum temperature differences (ΔT between center and edges) are calculated between -120 to -150°C (the glassy region).

As a final proof of principle, we nanowarmed a higher concentration of IONP (100 mgFe/mL EMG308) using VMP as the CPA (closely related to M22) in a 1mL cryovial [29]. We achieved warming rates at ∼1,500°C/min (Fig. S21). This is the fastest nanowarming rate that has been measured that we are aware of, supporting prior observations that rewarming rates will scale linearly with IONP concentration [28, 30, 31], which is promising for future development.

## Discussion

As a useful first-order approximation, the minimum cooling rates to avoid ice formation during the vitrification of different human organs can be predicted based on their characteristic length (L_C_) (Fig. 8A, Table S6). This can be used to establish CPA concentration and cooling protocol requirements (Fig. 8B, see supplementary for calculating min. vitrifiable CPA concentration). For instance, a small organ such as an ovary can convectively cool at ∼15°C/min, whereas the center of a liver is only expected to cool at ∼0.6°C/min (these calculations assume the organ is vitrified alone without any surrounding CPA, but L_C_ could be adjusted for individual container geometries). Note that all predicted cooling rates are greater than those achieved experimentally in the successfully vitrified 3L volume of M22 (cooling at ∼0.45°C/min).

**Fig. 8:**
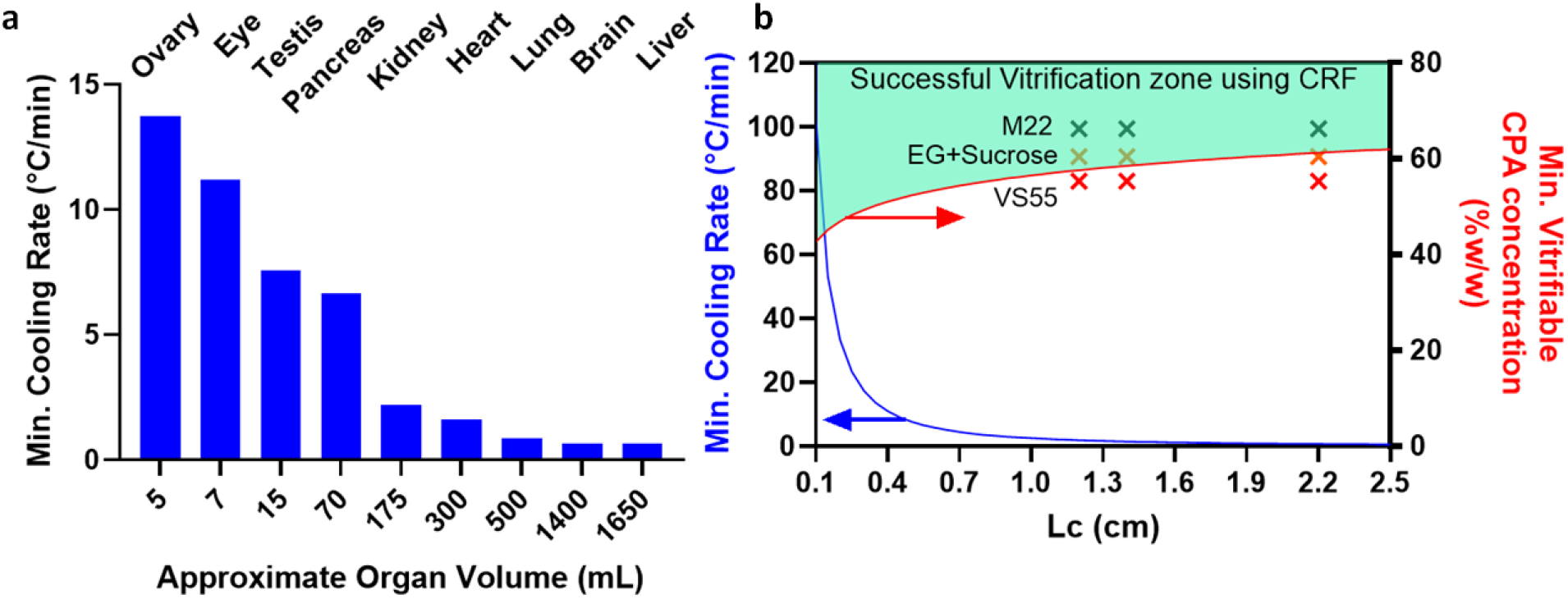
Achievable cooling rates and necessary CPA concentrations for human organ vitrification. **a** Predicted minimum cooling rate (CR) as a function of volume of various human organs. **b** Plot of minimum cooling rate vs. L_C_ (blue curve) and minimum vitrifiable CPA concentration in CRF as a function of L_C_ (red curve). L_C_ is calculated for various organs assuming ellipsoidal shape, and minimum cooing rate occurs at the center of geometry. L_C_, CR, and dimensions of organs can be found in Table S6. Note that to predict cooling rates at different volumes, one should calculate L_C_ and then use the blue plot in B. The experimental test points (x) showing success of M22, EG+Sucrose, and failure of VS55 at various L_C_ (0.5, 1, 3L volumes) are also plotted in B, validating the first-order approximation.

For this study, we chose CPAs with low CCRs (<1°C/min) [29, 32]. However, during CPA perfusion loading of organs, successfully perfused tissues may only equilibrate to ∼92-94% concentration of full-strength CPA [14, 33], which will substantially increase the required cooling rates. Thus, a conservative approach would be to select a CPA where the CCR at ∼94% loading is lower than the achievable cooling rate for the volume. For instance, using Fig. 8B, for 3 L we can calculate L_C_ ∼ 2.2 cm and the expected CR in the CRF to be < 0.8°C/min. Hence, the minimum CPA concentration for vitrification would be ∼62% w/w, which is slightly lower than M22 (∼66%w/w which includes carrier solution), where we have shown successful vitrification at 3L. Higher concentrations of CPAs such as VS83 (83% w/w CPA) have even lower CCR and can be more easily vitrified but increase biological toxicity relative to the CPAs chosen here [34]. To remain at a lower concentration of CPA and still achieve vitrification at higher volumes without toxicity, future work can assess the impact of ice recrystallization inhibitors (IRIs), polymers (e.g., polyglycerol-PGL, polyvinyl alcohol-PVA, polyethylene glycol-PEG, x-1000, z-1000, etc.), or other novel cryoprotective agents [35, 36]. Furthermore, due to larger heat transfer coefficients, convective cooling with liquid cryogens versus gaseous flow in CRF can enhance cooling rates [23].

The other major mode of physical injury during organ vitrification is cracking. To address this, it is crucial to thermally equilibrate samples through annealing, thereby reducing thermal gradients and stress before entering the glassy phase. Without proper annealing, these samples will be more susceptible to fracture during cryogenic storage and rewarming. Once annealed above T_g_ (where the supercooled CPA is fluid enough to relieve stresses), it is also critical to cool slowly enough to the storage temperature to avoid re-introducing significant thermal gradients and stress when cooling through the glass transition. To leverage these results for broader use and develop effective CRF cooling protocols (choosing annealing time, CR in the glassy region, equilibration time at storage, etc.) for volumes other than the ones analyzed in this study, we provide recommended annealing and glassy region CR as a function of L_C_ as plotted in Fig. S4. These protocols were designed to be faster than the CCR of the noted CPAs between the melt and glass transitions. Further, the ∆T within the glassy state was designed to be less than 20 °C, to avoid fracture. For larger volumes, the extended cooling protocols (i.e., ∼12 hrs for 3L) highlight the need for innovation, perhaps through a combination of convective and volumetric cooling approaches previously attempted in the kidney [37] and intestine [38].

While this study performed a careful analysis of cooling protocols necessary to achieve vitrified organ storage, another important factor that will be critical to organ banking is the effect of the long-term storage temperature and conditions. The storage temperature should be sufficiently below T_g_ to avoid the danger of devitrification, whereas storing at a temperature far below T_g_ may also induce more brittleness inside the samples [24]. For instance, in one study, storage of VS55 for 6 months just below T_g_ increased CWR from 50 to ∼100°C/min [39], suggesting additional ice nucleation may have occurred during storage. For long-term storage (days to months to years) of human organ-scale vitrified samples, physical aging of the glass can also elevate T_g_, decrease T_d_ (devitrification temperature), and increase heat release at the glass transition during rewarming from the vitrified state [40, 41]. This highlights the need for further study to identify the impacts of long-term vitrified storage and to identify optimal storage conditions. Further, CPA constituents (polymers, sugars, salts, etc.) can also impact stability in the brittle (i.e., glassy) state. For example, the thermal expansion coefficient of M22 (∼2.52 * 10^-4) is greater than VS55 (∼1.84 * 10^ -4) [42], which could mean that M22 (which has polymers that may be more thermally expansive [43]) might be more prone to fractures from thermal stress.

We were able to demonstrate uniform volumetric rewarming using nanowarming in volumes up to 2L. One hypothesized source of non-uniformity in larger RF coils is eddy current heating. However, prior analysis has assumed electrical properties at room temperature when in fact, eddy current heating is expected to be insignificant at cryogenic temperatures, especially in the vitrified glassy state (see supplementary eddy heating section and Fig. S19).

While we have shown that nanowarming provides uniform and scalable heating across various volumes, larger organ systems will likely achieve the greatest benefits. Fig. 9 (and Table S7) summarizes estimated nanowarming and convective rewarming rates for clinically relevant organs. While nanowarming offers substantial increases in rates and uniformity of heating for large organs, achievable convective rates may be higher (∼ 2 times) for small, less vascularized organs (e.g., ovaries and testes). However, a lack of uniformity during convective rewarming may still lead to cracking in these cases.

**Fig. 9:**
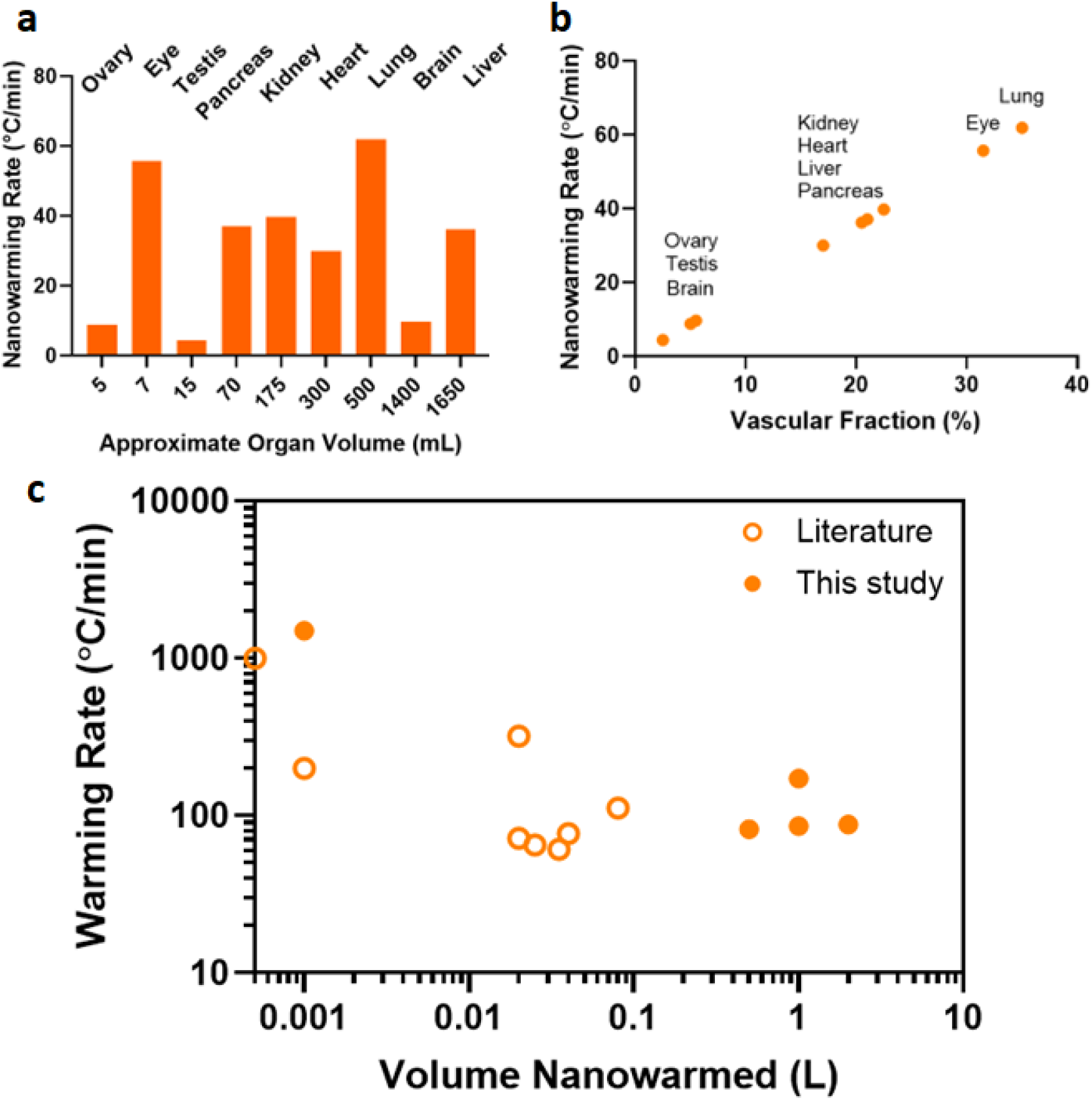
Nanowarming rate predictions for human organs. **a** Predicted nanowarming rate vs. total organ volume perfused at 10mgFe/ml. **b** Predicted nanowarming rates vs. vascular volume of various organs perfused with 10 mgFe/ml. Rates are estimated for 10mgFe/mL perfused IONP concentration with SAR_Fe_∼1050 W/gFe (see supplementary section for calculations and Table S7). **c** Nanowarming Rates for volumes from mL to L. Open circles represent literature values of warming rates achieved using nanowarming (see Table S8 for data values and references).

The CPA concentration needed for physical vitrification at 0.5L (relevant for human kidneys and hearts) is at least ∼8M (40%EG+0.6Msucrose) and increases to ∼9.4M (M22) at 3L scale (relevant for human livers) as the minimum expected cooling rates are below 1 °C/min (Table S6). To achieve warming rates above the CWR of those CPAs, the required IONP concentration and the SAR of those IONP should be carefully assessed. For instance, Fig. 9 shows that for a human kidney, perfused IONP concentration of ∼10 mgFe/mL will provide a rewarming rate of ∼40°C/min, which is safely above the CWR of M22. Fig. 9 also shows that highly vascularized organs such as lungs (non-alveolar portions) and eyes will produce significantly higher nanowarming rates (∼60°C/min) than convection alone (see Table S7 for convective warming rates). Note that the predicted nanowarming rate for adult human kidneys is lower than previously reported rates achieved in rat kidneys (40°C/min versus >60°C/min) [14]. In this case, it is anticipated that the rates achieved in the small volume of rat kidneys included contributions from ambient warming and a higher IONP concentration in the surrounding CPA than was achieved solely due to the vascular fraction of the rat kidney.

Higher rewarming rates will also be possible at higher IONP concentrations or with higher heating nanoparticles. We heated 100 mg Fe/ml IONP in VMP and reached rates up to ∼1,500°C/min (Fig. S21 and Fig. 9C). Notably, consistent with earlier studies, it is also interesting that we find higher heating and SAR_Fe_ in the cryogenic regime, likely due to the higher susceptibility and magnetization at such low temperatures and the reduced specific heat of CPAs in the glassy region [44]. It should be highlighted, too, that the above analysis is focused on the IONPs tested here and that other nanomaterials, such as nanowires/nanobars, have also shown ultra-rapid nanowarming heating rates up to ∼1,000°C/min at much lower nanoparticle concentrations in proof of principle cryovial (<1mL) samples (e.g., Cobalt Nickel nanowires in VS55) [45]. Still, these may be limited to external heating due to their aspect ratio, size, and concerns regarding biocompatibility.

Various other logistical factors must be considered to achieve human organ vitrification and nanowarming, including containers and IONP quantities. Verifying the cryogenic compatibility of the containers (i.e., polyimide, Teflon-PTFE, or other suitable material) and the ability to vacuum seal to remove the air interface as a nucleation site is key [46, 47]. Further, we can estimate the IONP quantity requirements for nanowarming a whole human organ. Using a human kidney as an example, the amount of IONP can be estimated as ∼ 0.4 gFe = 0.2 (vascular fraction) * 200 mL (organ volume) * 10 mgFe/mL (perfused IONP concentration). This amount should be multiplied by ∼2 to 4 times the vascular volume to ensure complete perfusion loading [13, 14], elevating the IONP required for a single kidney to ∼0.8-1.6 gFe. IONP will also be needed for the surrounding solution. This highlights the need for scalable, biocompatible, colloidally stable IONP production [30].

This study is limited by the assumption that CPA will fully equilibrate in tissue, but, as noted, minimum tissue equilibration in the range of ∼92-94% may be expected during organ perfusion [14, 33]. However, the analysis here can be adjusted to account for the expected dilution in CPA concentration, and as noted, these results suggest that vitrification is still achievable at these concentrations. Further, this work is limited to physical success, and considerations regarding the toxicity of specific CPAs in specific organs still need to be addressed. They will be critical in achieving successful organ banking.

## Methods

### Preparation of CPAs and Iron-oxide nanoparticle solutions

Three common vitrification CPAs were used in this study: M22, VS55, and 40%v/v EG+0.6M Sucrose, prepared according to their formulations and composition as listed in Table S2. The preparation of the CPAs was completed by weight-volume percent (M22, VS55) and volume-volume percent (EG+sucrose) using a volumetric flask, as previously reported [13, 18]. These CPAs were chosen due to their recent study in multiple organ systems, such as VS55 in rat hearts and kidneys [13, 15-17], M22 in rabbit kidneys [33], and 40%v/v EG+0.6M Sucrose in rat livers [18]. Commercially available iron-oxide nanoparticles (IONPs) EMG308 (Ferrotec, Bedford, USA) (∼10 nm) and silica-coated iron-oxide nanoparticles (sIONPs) [30] were used to prepare CPA + IONP mixture solutions. EMG308 was suspended in M22 with a modified carrier solution (water instead of LM5) to ensure the stability of these IONPs. sIONPs were synthesized as previously reported [30]. The colloidal stability of sIONPs and EMG308 in M22 was monitored by dynamic light scattering (DLS, see Fig. S1). The sIONPs and EMG308 were prepared in CPA (M22, VMP) or water. Recent work in organ vitrification was conducted with sIONPs [30, 48], but uncoated EMG308 was used for simplicity in some physical demonstrations in these studies.

### Heat Transfer Finite Element Modeling (FEM)

Computational modeling was performed using commercial multiphysics simulation software (COMSOL 5.4). A 3D CAD geometry of a cryobag filled with CPA was created and simulated in COMSOL for heat transfer simulations (Fig. S2). Three cryobag volumes were simulated (0.5, 1L, and 3L) with dimensions close to experimental volumes (see supplementary on cryobag finite element modeling (FEM)). Details regarding governing equations, boundary conditions, initial conditions, and geometry are provided in the supplementary text and Table S4. All the experimental and modeled cooling and warming rates are reported in the 0 to -100°C temperature range.

### Vitrification Experiments

Three volumes, 0.5, 1, and 3L, were evaluated for vitrification success. Polyethylene plastic “cryobags” of 2 mm thickness (McMaster Carr, Elmhurst, USA) were used to contain the CPA and organ samples (See Table S3 for bag sizes). Cryobags were heat-sealed at the top, and a clip was placed at the top to minimize air pockets and ice nucleation at the CPA-air interface. A controlled rate freezer (CRF) (Planar Kryo 560-II, Planar, Middlesex, UK) was used to execute cooling protocols for vitrification. Vitrified samples were stored in a -150°C cryogenic mechanical freezer (MDF-C2156VANC-PA, Panasonic, IL, USA). Physical vitrification was verified by transparency observed by visual inspection and photography. Temperature measurements were conducted using fiber optic temperature probes and a four-channel monitoring system (Qualitrol, Fairport, NY, and Micronor Sensors, Ventura, CA, USA) at 1-2-second intervals using FO Temp Assistant Software or Optilink Software (provided by the manufacturer). The probes were pre-calibrated for any offset in the cryogenic temperature range. Probe placement inside the cryobag filled with CPAs is shown in Fig. S11.

### Porcine Liver Perfusion and Vitrification

The University of Minnesota’s Institutional Animal Care and Use Committee (IACUC) approved this study. Porcine livers were recovered from male, 20-30kg, Yorkshire pigs sourced from a local vendor (Midwest Research Swine, MN). Animals were injected intravenously with 300 IU/kg heparin prior to euthanasia. After euthanasia, the abdomen was rapidly opened, the abdominal aorta and portal vein were cannulated, the thoracic aorta was cross-clamped, the suprahepatic vena cava (SHVC) was vented, and organs were flushed immediately with 5L of a cold Histidine-Tryptophan-Ketoglutarate (HTK) solution. Once the flush was complete, the liver was explanted and placed in a cold HTK solution for transport. After back table preparation, the porcine liver was perfused via the portal vein with 40%EG+0.6M Sucrose in a step-loading protocol similar to that published for rat livers [18]. Specifically, 10%EG and 25%EG were perfused for 15 minutes each, followed by 40%EG+0.6M sucrose for 70 minutes at a constant flow rate of ∼65 mL/min (see Fig. S5). More details can be found in the supplementary information.

### μCT Imaging for verification of vitrification

Microcomputed tomography (μCT) was used to verify physical vitrification. A foam/plastic container was used to hold the 0.5L samples during μCT scanning, similar to that used previously for 20 mL systems [49]. The samples were scanned in a μCT imaging system NIKON XT H 225 (Nikon Metrology, MI, USA) for 500mL volumes. Detailed information about μCT settings and image reconstruction can be found in supplementary information.

### RF Coil Characterization (120kW)

AMF LifeSystems, LLC. (Auburn Hills, MI) designed and built a custom-RF coil specifically for the development of nanowarming. Performance was characterized by measuring the spatial distribution across the full range of magnetic field values. To determine magnetic field strength (axial and radial component), we used a 2D high-frequency RF probe (AMF LifeSystems) placed inside the RF coils. We used an oscilloscope (LA354, LeCroy, NY) to record the voltage signals, which were then converted to magnetic field values. We used modeled as well as experimentally measured magnetic field distribution to compare spatial uniformity between the previously characterized 15 kW [6] and the new 120 kW RF coils (Fig. S15). The supplementary materials provide a detailed description of the RF coil in Figs. S15-S17 and Table S3.

### SAR Measurements

Specific absorption rates (SAR) measurements were performed on a 1mL sample volume of CPA with iron-oxide nanoparticles (EMG-308, sIONPs at ∼4mgFe/mL) in a cryovial. The cryovial was placed inside the RF coils (15, 120kW systems) within an insulated 3D-printed holder, and the temperature was recorded using fiber optic temperature probes. SAR was then calculated using the time rise method described elsewhere [25, 50] (see supplemental SAR section, Fig. S18).

### Bulk Nanowarming Experiments

Liter-scale nanowarming was performed in the 120 kW RF coil described above. Volumes of 0.5, 1, and 2L M22 with EMG308 were vitrified in heat-sealed cryobags with cylindrical shapes that fit within the RF coil. Three fiber optic temperature probes were placed in the center, left, and right regions of the cryobag ∼4cm apart for all volumes using 3D printed jigs (Fig. S20).

After vitrification, samples were stored overnight at -150°C in a cryogenic freezer (MDF-C2156VANC-PA, Panasonic, IL). For rewarming, the sample was rapidly transferred from the freezer into the 120k kW RF coil (< 20 seconds), placed in an insulated holder, and immediately rewarmed. The temperature was recorded every 1 sec during rewarming using the fiber optic thermometry system described above.

### Data analysis

All the graphs in the figures are plotted using GraphPad Prism 10. A total of ≥3 replicates are conducted for all the cooling, nanowarming, and SAR experiments. The number of replicates is listed in each figure legend.

## Acknowledgment

This work was supported by the National Institutes of Health (NIH) Grant R01DK117425, NIH Grant R01HL135046, NIH Grant R01DK132211, and National Science Foundation (NSF) Grant EEC 1941543. We acknowledge support from a gift from the Biostasis Research Institute, funded partly through contributions from LifeGift, Nevada Donor Network, LifeSource, Donor Network West, and Lifebanc. The authors would like to thank the Dental CT laboratory for μCT scans of CPA volumes and the University of Minnesota, Minnesota Supercomputing Institute (MSI) for computational FEM access. We acknowledge Elliott Magnuson for the initial work on cryobags, Mikaela Hintz for technical assistance with CPA preparation, and Andy Grams for graphical design illustration. Lastly, we would also like to acknowledge Dr. Anirudh Sharma for his initial scientific work on RF coil scaling and Dr. Rhonda Franklin for scientific discussion on RF coil eddy heating.

## Ethical Declaration

This study complies with all relevant ethical considerations. The Institutional IACUC committee from the University of Minnesota (protocol #2111A39564) approved all animal studies.

## Data Availability Statement

All the data supporting this study’s findings are available in the supplementary material. A source data file is also provided with this article. Any assistance in accessing or interpreting any data can be requested by contacting J.C.B., corresponding author

## Author’s Contributions

L.G. conceived and performed the study as a whole and contributed to all aspects with support and supervision from J.C.B. and E.B.F. L.G. carried out all experiments involving CPA, CPA+IONPs, porcine liver vitrification, and nanowarming with input from B.E.N., Z.H., and M.L.E. L.G. performed computational heat transfer modeling and SAR theoretical modeling with input from M.L.E. B.E.N performed liver procurement surgeries and provided technical assistance in liver perfusion experiments. C.S. prepared all the CPA and IONP solutions and conducted stability studies. S.K. conducted relaxometry and ICP-OES for iron quantification. Z.H. performed μCT on all CPA samples with assistance from S.K. L.G. conducted all the SAR measurements with some measurements performed by M.L.E. L.G. performed 120 kW RF coil characterization with assistance from R.G. R.G. performed the modeling of magnetic field distribution in RF coils. L.G. did all the data analysis. L.G. wrote the manuscript with input from M.L.E., E.B.F, and J.C.B. All authors reviewed, edited, and provided critical feedback. J.C.B. and E.B.F. acquired funding, and J.C.B. oversaw project administration.

