## Supplementary Information for "Physical vitrification and nanowarming at human organ scale to enable cryopreservation"

**Table of Contents**

***Supplementary Method, Results and Discussion***

ICP-OES quantification of IONP solutions……………………………………………………………..3

Computational modeling of heat transfer during vitrification……………………………………….3

Porcine liver perfusion………………………………………………………………………..…………6

Porcine liver heat transfer modeling………………………………………………………………….7

Detailed results for rooling CPA cryobags in CRF………………………………………………….9

µCT imaging to verify vitrification of cryobags…………………………………………………...…..12

120 kW RF coil characterization……………………………………………………………………...14

***Supplementary Figures***

Fig. S1: Stability of EMG308 in M22 by DLS with and without carrier solution (LM5)…………….3

Fig. S2: Cryobag heat transfer model…………………………………………………………………..4

Fig. S3: Cryobag model thermal response…………………………………………………………….5

Fig. S4: Designing CRF cooling protocols based upon L_C_…………………………………………...6

Fig. S5: Perfusion of a ~0.8L porcine liver with 40%EG+0.6MSucrose…………………………….7

Fig. S6: Model prediction of porcine liver vitrification in CRF. ……………………………………….8

Fig. S7: Placement of same sample volume, i.e., 1 L, in 3 different orientations inside CRF resulting in three unique characteristics length………………………………………………………..8

Fig. S8: Representative photos of ice formation failure in cryobags………………………………..9

Fig. S9: Representative photos of fracture failure in cryobags………………………………………9

Fig. S10: Photos after executing CRF cooling protocols for the three CPAs (M22, 40% EG+0.6M Sucrose, VS55) tested at 0.5 L, 1L and 3L…………………………………………………………...10

Fig. S21: Nanowarming heating demonstration at multiple scales (mL to L)……………………..21

***Supplementary Tables***

Table S1: Scale of systems successfully vitrified and rewarmed……………….………………….24

Table S2: CPA composition and calorimetric properties…………………………………………….25

Table S3 Details of cryobag dimensions used in the liter-scale vitrification and nanowarming study………………………………………………………………………………………………………26

Table S4: Thermo-physical properties of CPA used for computational FEM modeling………….26

Table S5: Capabilities of custom-built state-of-art 120kW RF Coil System……………………….27

Table S6: Center cooling rates for human organs (no surrounding solution or container) based on characteristic lengths………………………………………………………………………………..27

Table S7: Nanowarming rates for organs loaded with 10mgFe/mL IONPs…………………….....28

Table S8: Nanowarming rates reported in cryopreserved volumes prior to this study…………..29

***Supplementary References***......................................................................................................30

ICP-OES quantification of IONP solutions

Total Fe content (used to calculate sIONP concentration) in solutions of EMG308 in M22 with LM5 removed and sIONPs in M22 were measured using ICP-OES by pipetting 20 µL of each solution into a glass ampule containing 400 µL HNO3 and 180 µL DI water for a total solution volume of 600 µL. The ampules were flame-sealed, and the solution was digested overnight at 90°C. Solutions were diluted 10x in DI water before total Fe measurement via ICP-OES (University of Minnesota Research Analytical Laboratory, St. Paul, MN).

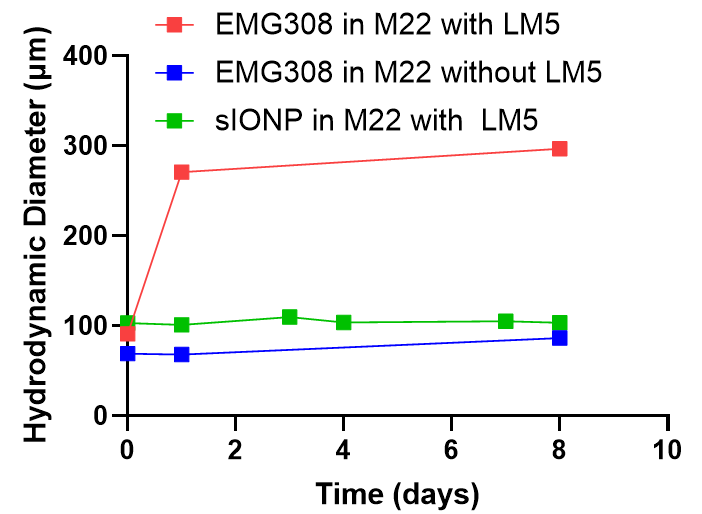

**Fig. S1: Stability of EMG308 in M22 by DLS with and without carrier solution (LM5).** An increase in hydrodynamic diameter corresponds to aggregation and instability of EMG308s when carrier solution is present (LM5 in M22). sIONPs are stable in M22 even with carrier solutions (LM5).

Computational Modeling of Heat Transfer During Vitrification

The governing heat transfer equation is:

$\rho C_{P}\frac{\partial T}{\partial t}=\nabla.(k\nabla T){+q}_{v}^{'''}$	(1)

where T is temperature, ρ is density, C_P_ is specific heat capacity, k is thermal conductivity, and q_v_’’’ is the heat generation term. Details of mathematical modeling are given in Fig. S2. A cryobag with CPA was modeled as a simplified geometry assuming a height equal to the actual height of CPA during experiments. The CAD geometry was created in COMSOL 5.4, and meshing was performed using the default setting of extra fine tetrahedral mesh (maximum element size ~ 5mm). The dimensions (Width x Height x Thickness) of the modeled cryobag for 0.5L volumes were 14 x 13 x 5.5 (L_C_~1.19cm), 1L were 19.5 x 14.5 x 6.5cm (L_C_~1.42cm), and 3L were 18.5 x 30 x 10.5cm (L_C_~2.19 cm). Fig. S2 shows the computational geometry of the cryobag with CPA for 0.5, 1.0, and 2.5L volumes. Temperature is plotted for the center of the cryobag geometry (geometric center with slowest response) vs. the temperature at the edge of the cryobag (assumed 95% of half-width from the center and reflective of fastest response). The precise edge would be less informative as it follows the boundary condition (e.g., CRF temperature in cooling). The average center Cooling rate (CR) is estimated by calculating instantaneous dT/dt and taking its average from 0 to -100°C range. Thermal properties are assumed to be that of the CPA M22 for all the volumes analyzed (see Table S3). Some CPAs, such as VS55, have lower specific heat (by ~10%), which would lead to faster CRs than conservatively predicted here.

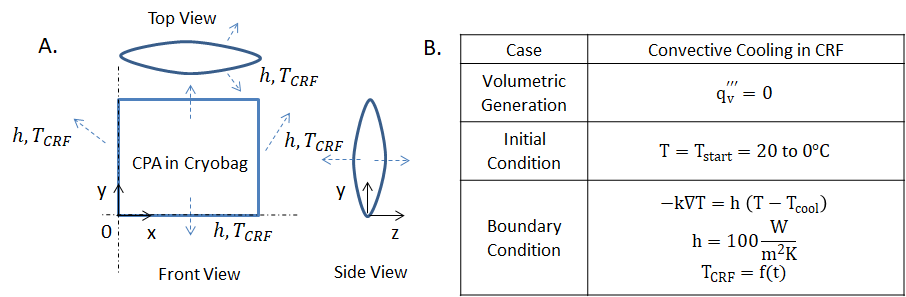

**Fig. S2: Cryobag heat transfer model.** A. Schematic of the geometry, and B. Table with volumetric heat generation, initial and boundary conditions. h = 100 W/m^2^K is assumed in the control rate freezer (CRF) based upon literature modeled fitting of experimental temperature [10, 24].

The maximum temperature difference across the sample to avoid fracture (i.e., predicted that the sample will fracture above this threshold) can be estimated from a simple thermal shock equation below.

$\Delta T_{\mathrm{limit}}=\sigma_{\mathrm{tensile}} \frac{(1-v)}{gE\beta}$	$\Delta T_{max}=\sigma_{tensile} \frac{(1-v)}{gE\beta}$	(2)

Where g is the geometric coefficient (assumed as 0.5 for cylindrical geometry), ν is Poisson’s ratio (assumed as 0.2 for typical brittle materials), E is the modulus of elasticity (assumed as 1 GPa for organic materials), and σ is the tensile yield strength of CPA (adapted as 3.2 MPa) based upon prior literature [30]. Therefore, a conservative estimate of ΔT_max_ threshold can be estimated for M22 by assuming the largest coefficient of linear thermal expansion: β (~2.52*10^-4 [1/K]) [31], which comes out to be ~20°C.

Note that the actual threshold from the thermal shock equation would be larger as β is a function of temperature and would decrease at cryogenic temperatures [31]. Furthermore, for other CPAs, such as VS55, the thermal expansion coefficient (β) has been reported to be slightly smaller (~1.84 * 10^-4), implying the ΔT_max_ threshold would be higher. This threshold from the thermal shock equation is a simplified estimate with limitations such as assumed material properties and lack of length scale correlation and, hence, should be validated experimentally. Other criteria for indirectly estimating a threshold for thermal stress could be used, such as based on cooling rate and characteristics length as given in the literature [32]. For two similar biomaterials, such as organs varying in size, CR*L_C_^2^ = constant, where CR is taken to be the outer surface cooling rate, and L_C_ is assumed to be the effective length scale of the outer surface. This allows a simple criterion in choosing the cooling rate in the glassy region to minimize thermal stresses below a threshold (smaller CR for a larger L_C_).

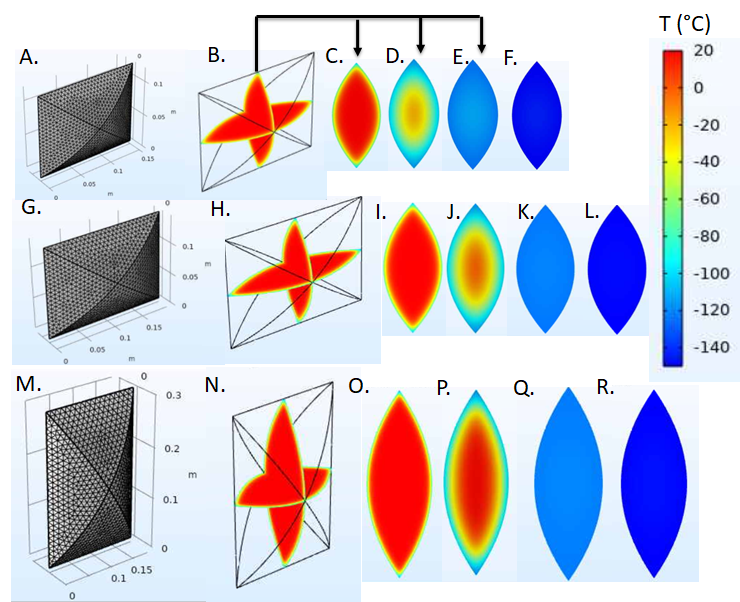

**Fig. S3: Cryobag model thermal response**. Computational mesh geometry for A. 0.5L, F. 1L, K. 3L volume. We modeled temperature distribution within the cryobag at two intersecting central planes for B. 0.5L, G. 1L, and L. 3L volume at a time ~ 5min (start of annealing step). We modeled temperature distribution in one full 2D plane within cryobag at the time points at the start of annealing (t = 5 min) C. 0.5L, H. 1L, M. 3L, 30 mins after start of annealing (t = 35 min) D. 0.5L, J. 1L, P. 3L, end of annealing D. 0.5L (t ~ 184 min), I. 1L (t ~ 254 min), N. 3L (t ~ 524 min) and end of cooling protocol E. 0.5L (t ~ 275 min), J. 1L (t ~ 374 min), O. 3L (t ~ 754 min) volume.

The effect of start temperature on CRF cooling protocol is also analyzed by varying the start temperature of the cryobag sample and CRF (20, 0, -20°C), as shown in Fig.S4. Generally, high starting temperatures lead to slightly faster average cooling rates (although the difference is marginal in the critical range for supercooling), but at the cost of longer annealing times. However, the start temperature cannot be higher than the organ perfusion temperature. For this reason, we chose 0°C. Characteristics length for heat transfer in a complex shape/geometry is calculated as L_C_ = V/A where V is total volume, V is the total volume, and A is the total surface area of the geometry participating in heat transfer. For designing CRF vitrification protocols other than the cryobag volumes tested in this study, we provide cooling protocol parameters as a function of L_C_, which will facilitate reasonable approximations for successful vitrification of different geometries (see Fig. S4 G).

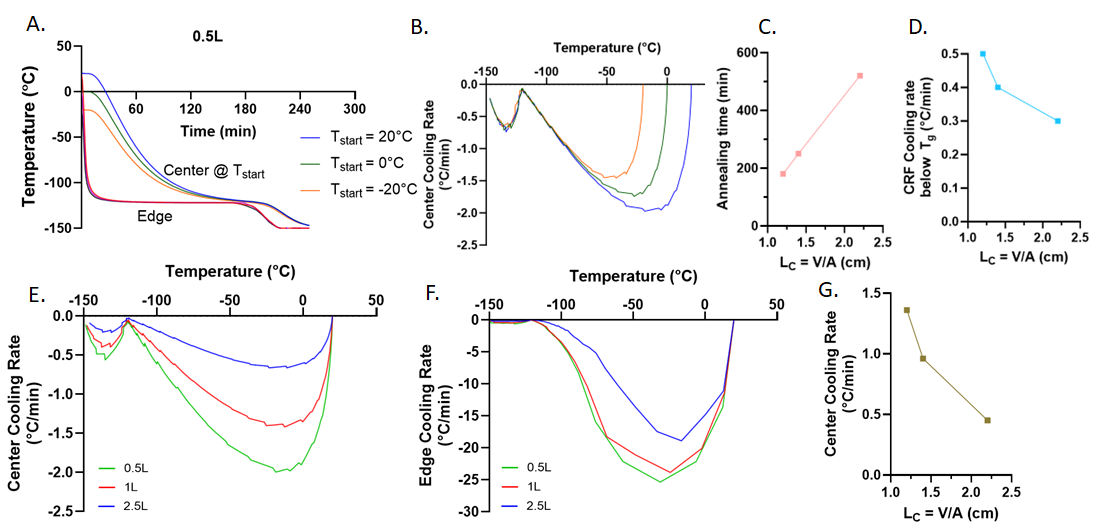

**Fig. S4: Designing CRF cooling protocols based upon L_C_.** A. Temperature vs. time for 0.5 L M22 (CPA) volume with a varying start temperature. B. Center cooling rate vs. time for varying start temperature for 0.5L M22. C. Plot of annealing time as a function of L_C_. D. Plot of CRF cooling rate needed for different L_C_ in the glassy region (from T_anneal_ to T_storage_). E. Center and F. Edge cooling rate variation with temperature for 0.5, 1, and 3L M22 (CPA) volumes. G. Average center cooling rate (0 to -100°C) as a function of L_C._

Porcine liver perfusion

Constant flow-rate machine perfusion of porcine livers was used to load EG-Sucrose based on a modified protocol, as [12] shown in Fig. S5. Step loading began with carrier solution Euro-Collins (EC) at ~65 mL/min. It was based on hypothermic machine perfusion [33] and supercooling studies [34], which maintained portal venous perfusion pressure at physiological values (<3-4 mmHg). For this study, we perfused CPA long enough to achieve >96% equilibration in tissue to demonstrate physical vitrification success (verified using the effluent reflective index). As in previous hypothermic machine perfusion studies, the temperature was kept close to ~4°C [12, 14]. Note that this protocol was adapted to provide a demonstration of physical success. No biological assessment of the organ was performed, and it is anticipated that further modification to this protocol will be required to minimize CPA toxicity and achieve biological success.

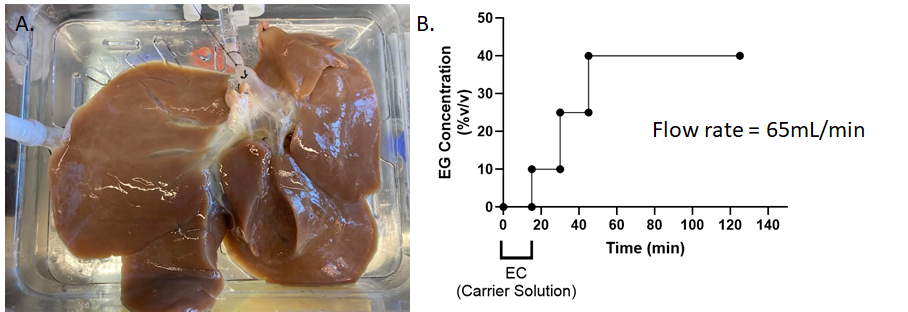

**Fig. S5: Perfusion of a ~0.8L porcine liver with 40%EG+0.6MSucrose**: **A** Photo of the liver after cannulation via the portal vein. **B** Step loading of CPA (concentration vs. time). Note that the first 15 minutes correspond to 0%EG (i.e., Euro Collins-carrier solution only).

Porcine liver heat transfer modeling

The ~0.8L liver in a cryobag was modeled as a system with dimensions: width x height x thickness ~ 20 x 30 x 2cm, as shown in Fig. S6. The CAD geometry was constructed from the actual photos of the liver (Fig. 4) in cryobag using SOLIDWORKS and later imported into COMSOL 5.4 for heat transfer simulation. Total volume was ~ 1L, laid flat in the CRF, thereby decreasing L_C_ to ~0.83cm, allowing execution of the “0.5L” (L_C_~1cm) CRF cooling protocol (See Fig. 3). Reducing the thickness of the sample inside the cryobag (i.e., reducing L_C_) increases the surface area for heat transfer and simultaneously reduces the length over which heat must be conducted. The predicted ΔT_max_ was ~ 3°C when transitioning to the glassy state, minimizing the risk of fracture failure (Fig. S6E). The modeled CR within the liver (Fig. S6F) at the center is ~4°C/min (averaged over 0 to -100°C), which is greater than the CCR of 40%EG+0.6MSucrose (<1°C/min). Note in experiments, the porcine liver (total bag volume ~1L= liver ~0.8L + surrounding CPA ~ 0.2L) leads to an L_C_ of ~0.8 cm (20x24x2cm), which is smaller than the L_C_ of an adult human liver (~2 cm) and resembles the size of a juvenile human liver or adult human kidney (L_C_~1.1 cm).

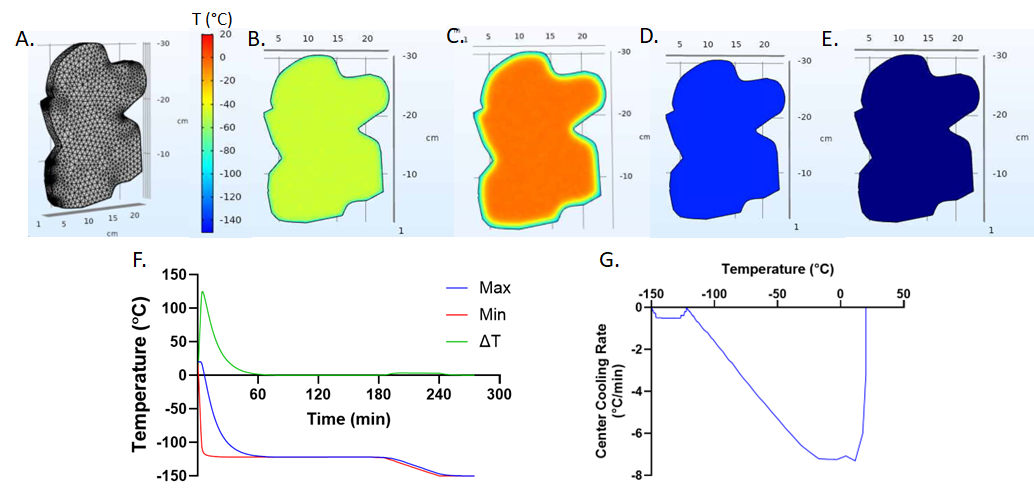

**Fig. S6: Model prediction of porcine liver vitrification in CRF.** A. 3D CAD geometry. B. Temperature distribution across the liver at the start of the annealing step (t ~ 5 min), C. 5 min after the start of annealing (t ~ 100 min) D. distribution at the end of the annealing (t ~ 184 min), and E. at the end of the full protocol (t ~ 275 min). F. Temperature vs time plot for liver. Max and Min represent the maximum and minimum temperatures evaluated over the completed domain volume. Note that the center of the liver corresponds to the slowest rate of cooling (“Max” temperature in the liver). The temperature difference between Max and Min is also plotted. G. Plot of center cooling rate with temperature (average ~ 4°C/min over 0 to -100°C).

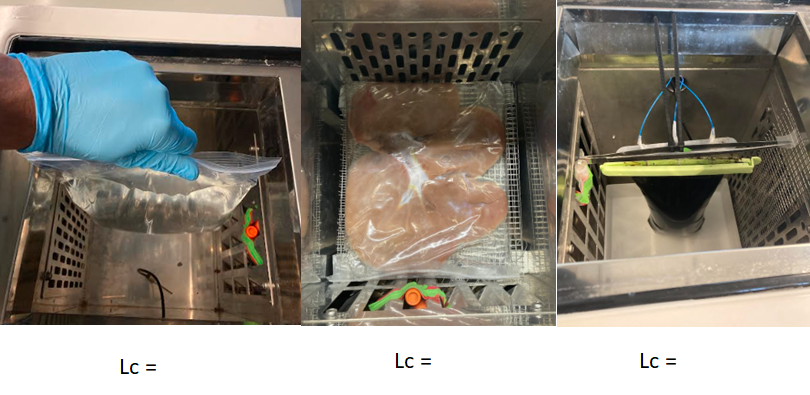

**Fig. S7: Placement of same sample volume, i.e., 1 L, in 3 different orientations inside CRF resulting in three unique characteristics length A. L_C_ ~ 1.4 cm, B. L_C_ ~ 1 cm, C. L_C_ ~1.5 cm.**

Detailed results for cooling CPA cryobags in CRF

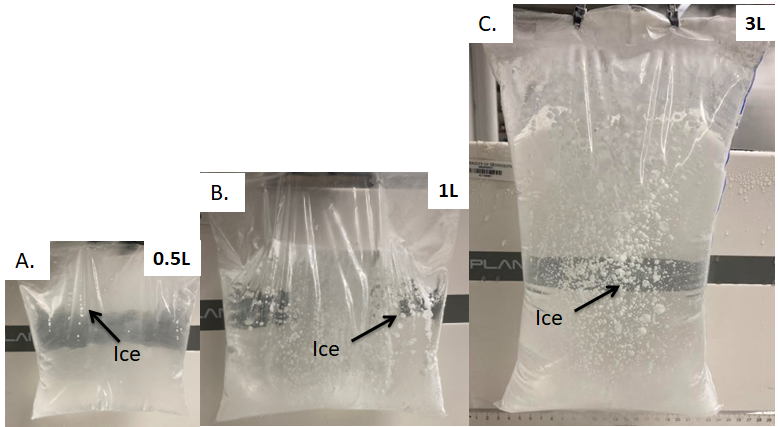

**Fig. S8: Representative photos of ice formation failure in cryobags filled with A. 0.5, B. 1, and C. 3 L volumes of CPA VS55.** Ice crystallization occurs if the sample cooling rate is less than the CCR of the CPA. Ice nucleation and growth depend on the sample's local temperature, which is a stochastic process, so note the formation of distributed ice spherulites.

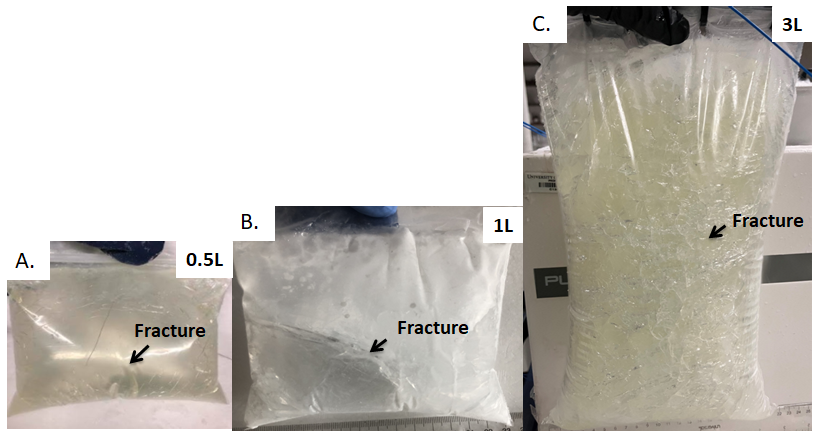

**Fig. S9: Representative photos of fracture failure in cryobags filled with A. 0.5, B. 1, and C. 3 L volumes of CPA M22**. These fractures occur due to thermal stress if the sample is not thermally equilibrated (annealed) above T_g_ or cooled too rapidly while in the glassy state.

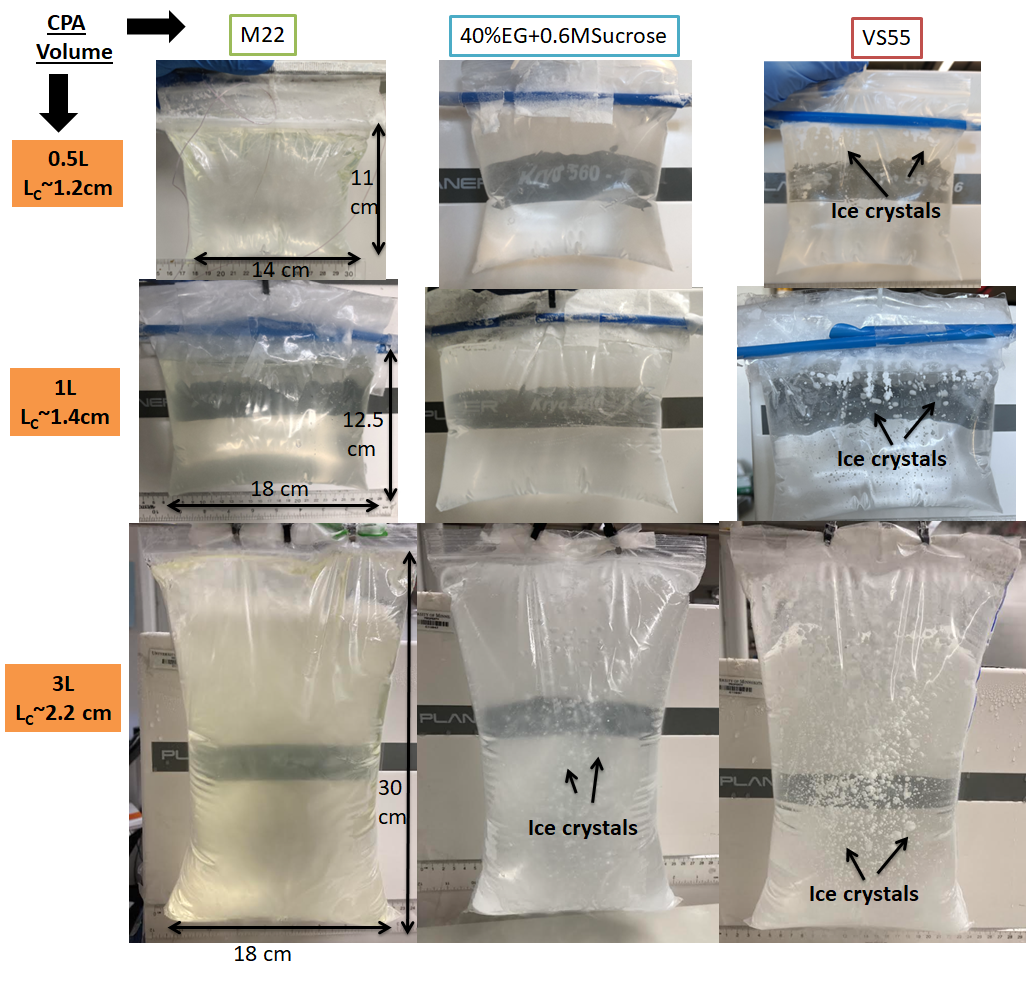

**Fig. S10: Photos after executing CRF cooling protocols (Fig. 3c in main text) for the three CPAs (M22-left, 40% EG+0.6M Sucrose-center, VS55-right) tested at 0.5 L (A, B & C), 1L (D, E, & F) and 3L (G, H & I) with L_c_’s noted**. VS55 forms ice at all volumes. 40%EG+0.6MSucrose successfully vitrified at 0.5 and 1L but not 3 L. M22 successfully vitrified at all three volumes (up to 3L). The CRF protocols were effective for cooling without fracture in all cases.

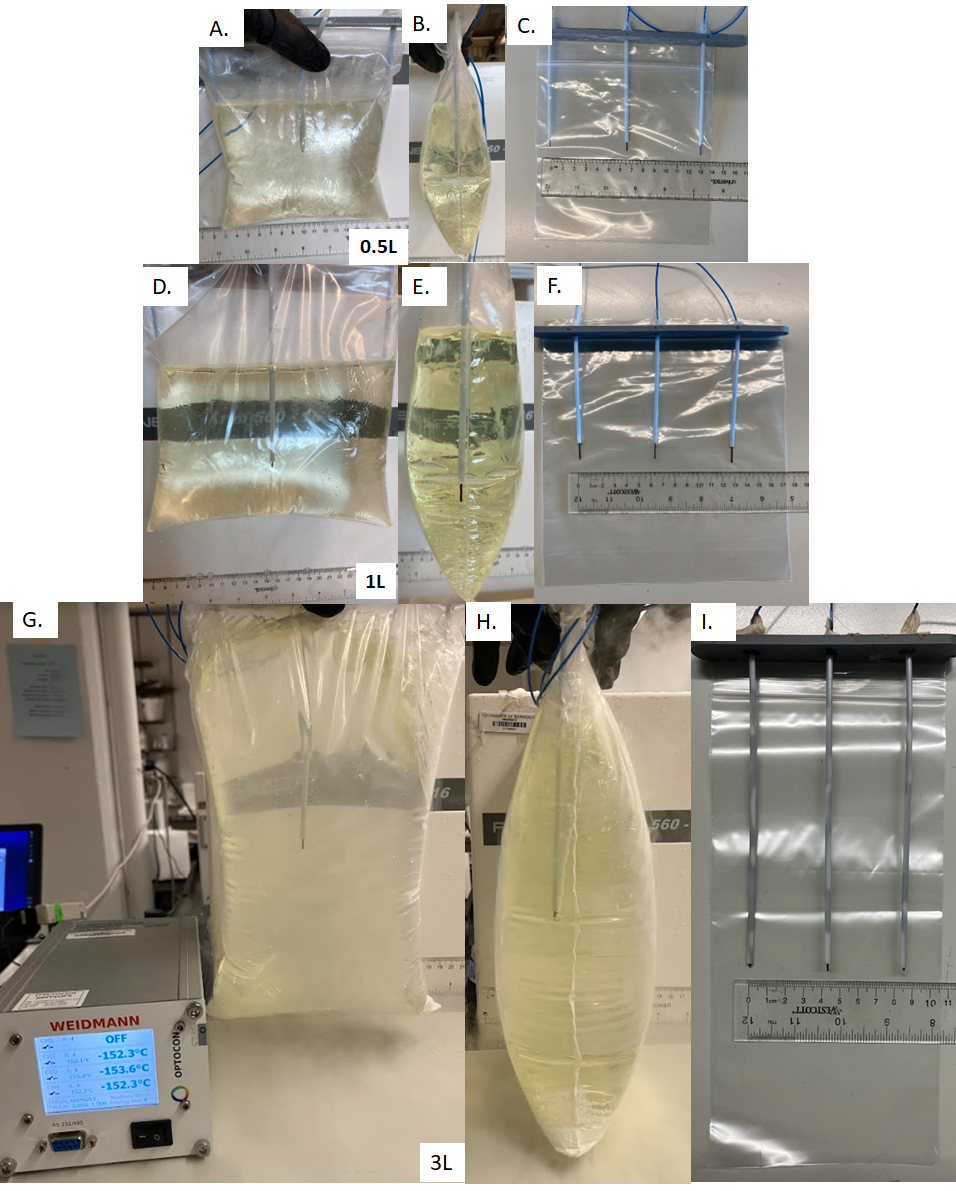

**Fig. S11: 3D printed jigs for fiber optic temperature probe placement in cryobag**. A. Front and B. side view of 0.5L M22 vitrified with probes. C. Probe location in 0.5L cryobag. D. Front and E. side view of 1L M22 vitrified with probes. F. Probe location in 1L cryobag. G. Front and H. side view of 3L M22 vitrified with probes. I. Probe location in 3L cryobag.

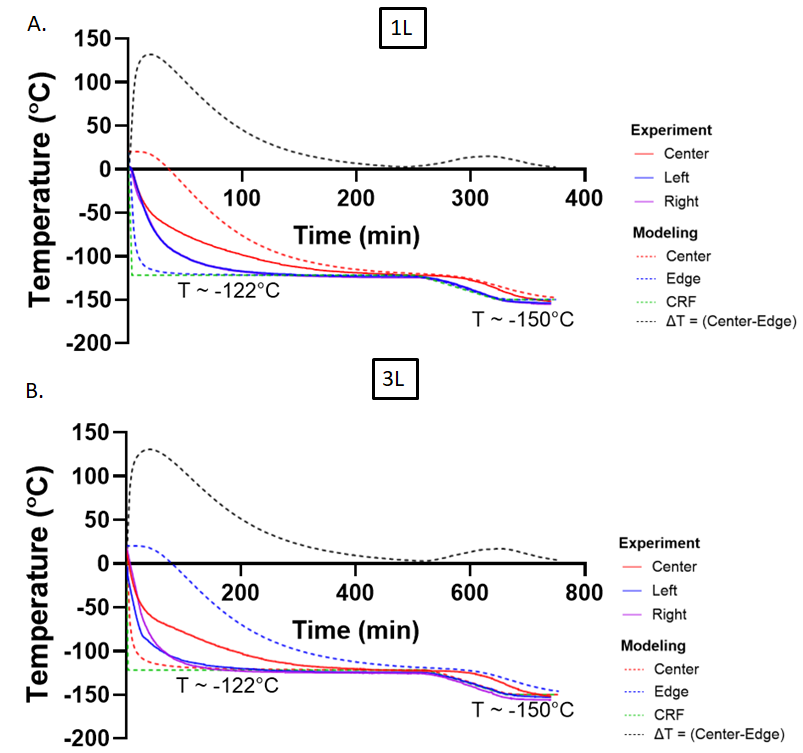

**Fig. S12: Temperature vs. time response of M22 cryobags during vitrification in CRF**. A. 1L and B. 3L. Solid lines represent experimentally measured temperature, and dashed lines are predictions from the heat transfer modeling. See Fig. 3 in the main text for the 0.5L temperature vs time plot.

µCT imaging to verify vitrification of cryobags

Cryobags were imaged using a µCT scanner (NIKON XT H 225) for 500 mL volumes. M22, VS55, and 40%EG+0.6Msucrose were vitrified in the CRF and transported to the µCT lab using an LN2-based transport container. Protocols and containers were modified based on previous work in rat organs [12, 13]. For the NIKON XT H 225 CPA scans, the accelerating voltage was set at 121 kV, and the current was set to 150 μA. We placed a 1-mm aluminum filter between the source and the object to reduce the beam-hardening effect [10]. The resolution was 0.098 mm for 0.5L samples. During imaging, the vitrified CPA samples were held in LN2 vapor at a temperature of approximately -150°C in a Styrofoam container. Two calibration samples, i.e., air and water at room temperature, were attached to the top of the vitrified CT sample container and used to calculate Hounsfield unit (HU) radiodensity. The samples remained in the glass state throughout the ~30-minute imaging time. Artifacts such as beam hardening are addressed during reconstruction to enhance image quality (3D CT pro, Nikon Metrology, MI). Lastly, image post-processing is executed using VGstudio Max 3.2 (Volume Graphics, NC, USA) to convert into unsigned 16-bit float images, which are later exported as DICOM images for a final analysis using MATLAB (MathWorks). HU are calculated based upon the following formula:

$HU= 1000*\frac{Grey scale Sample-Grey scale Water}{Grey scale Water- Grey scale Air}$ (3)

The edge of the sample shows a smaller signal (Fig. S13) due to common edge artifacts, such as the beam hardening effect, which we attempted to reduce but didn’t completely eliminate.

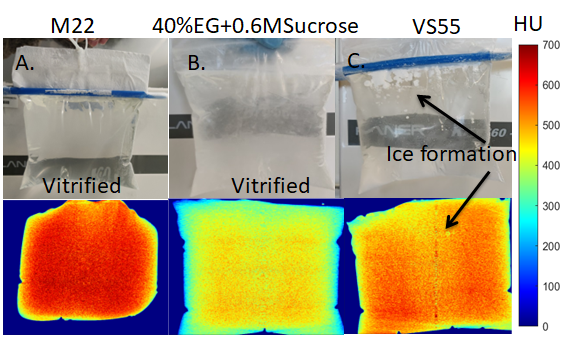

**Fig. S13: Photographs and µCT images of 0.5 L cryobags of M22, EG-Sucrose, and VS55 after cooling**. The pseudo-color sidebar shows radiodensity in Hounsfield units (HU). VS55 shows some ice formation, whereas M22 and 40%EG+0.6MSucrose show complete vitrification. Note that the HU for vitrification is different for 40%EG+0.6MSucrose (HU> 400) than M22 and VS55 (Vitrified HU > 500).

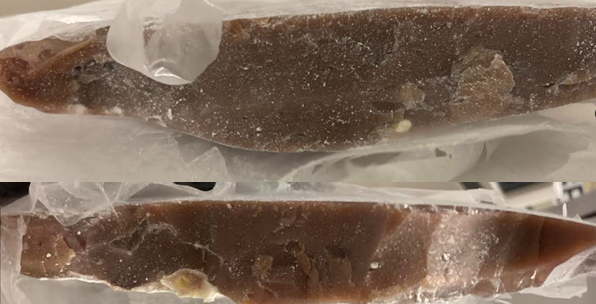

**Fig. S14: Photos of a center cross-section of vitrified bisected pig liver**.

120 kW RF coil characterization

Electromagnetic models for both systems were performed using Flux2D software (Altair), assuming rotational symmetry, frequency domain, and a voltage source.  This approach was previously validated for the 15 kW (80 mL) [16, 35] and 120 kW (2.5 L) coils [36]. The generated magnetic field variation within the coil volume is plotted in Fig. S15, with the 120 kW coil being the most uniform field in the largest volume of interest (VOI). For 120 kW RF coil spatial field characterization (shown in main Fig. 5c), axial location is varied by moving the probe along the axis of the cylindrical coil volume from 0 (inner end of coil sample space) to 20 cm (towards outer end of sample space). Radial location is varied by moving the probe radially into different holes of the jig (green rectangle box) spaced at 0, 2, and 4 cm apart.

A simple linear least square fit (equation 4) to magnetic field strength and coil voltage (Fig. S16) gives a correlation, which is useful in setting up a desired magnetic field strength for nanowarming experiments.

$H \left( \frac{kA}{m} \right)=0.02473 Voltage \left( V \right)$(4)

Magnetic field strength is calculated from voltage readings from the oscilloscope using the simple equation below:

$B_{axial}=\frac{V_{rms}}{s_{axial}*f}$ (5)

$B_{radial}=\frac{V_{rms}}{s_{radial}*f}$ (6)

$B_{total}=\sqrt{{B_{axial}}^{2}+{B_{radial}}^{2}}$ (7)

where B is magnetic flux density, V_rms_ is measured voltage in root mean square (RMS), *f* is frequency in kHz, and s_axial_ and s_radial_ are sensitivity factors for axial and radial direction (calibrated for each probe). Magnetic field strength (H) can be found by:

$H=\frac{B}{\mu_{o}}$ (8)

where µ_o_ is vacuum magnetic permeability.

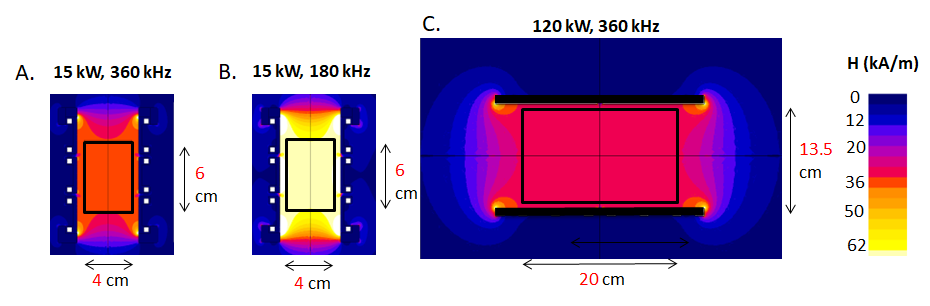

**Fig. S15: 2D FEA simulation of uniformity**. A. 15kW coil at peak field ~35 kA/m and 360 kHz B. 15kW coil at peak field ~60 kA/m and 180 kHz, C. 120kW coil at peak field ~ 35 kA/m and 360 kHz.

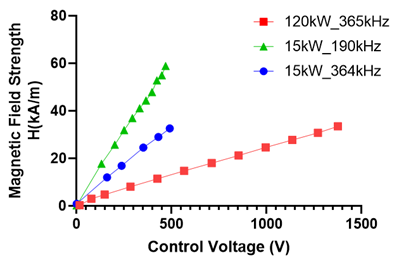

**Fig. S16: Measured magnetic field strength**. Field strength vs. control voltage for three RF coils 1kW, 15kW, and 120kW used for nanowarming. Frequency variation is within 365 ± 5kHz, 360 ± 4kHz and 190 ± 10kHz.

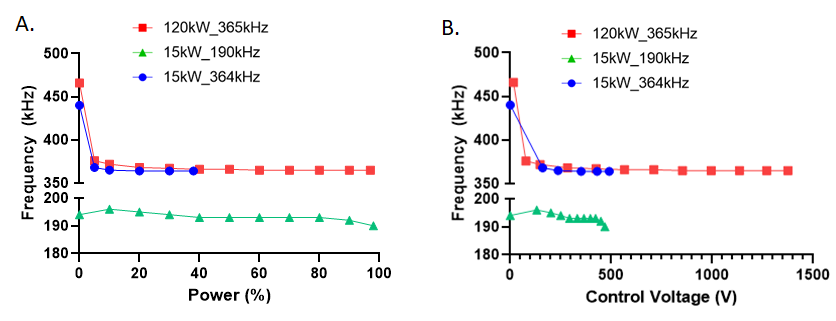

**Fig. S17: RF Coil frequency**. Frequency measured using an oscilloscope as a function of A. coil power and B. control voltage for the three RF coils 1, 15, 120kW systems.

Experimental SAR

SAR was measured based on previously published methods [37]. EMG308 IONPs were prepared at ~4mgFe/mL (ICP-OES: 4.38 mgFe/mL) concentrations in M22, and H20 while sIONP were prepared at ~4mgFe/mL (ICP-OES: 3.5 mgFe/mL) in M22. 1 mL of IONP solution in a cryovial was heated in an RF coil for ~60-180 sec or more depending on the achieved heating rate (Fig. S18A-C). Slower rates at lower field strengths were heated for a longer duration (for example, ~180 sec at 5 kA/m). Experimental SAR_V_ is directly calculated by measuring the temperature rise produced while heating IONP samples inside an RF coil and calculated (time-rise method) as previously reported [38] and shown briefly below:

${SAR}_{V}=\rho C_{P}\frac{\partial T}{\partial t}$	(9)

${SAR}_{Fe}*C_{Fe}= {SAR}_{V}$	(10)

where ρ is density (~997 kg/m^3^ for water, ~1100 kg/m^3^ for M22 samples), C_Fe_ is concentration of IONPs (~4.4 mgFe/mL for water, ~3.5 mgFe/mL for M22 samples), and C_P_ is specific heat at constant pressure (~4180 J/kg.K for water, ~3500 J/kg.K for M22 samples). SAR can be expressed as heat generation per unit volume in terms of SAR_V_ (W/m^3^) or heat generation per unit mass of magnetic nanoparticles (normalized to mass iron (Fe) based on standard convention), commonly called SAR_Fe_ (W/g.Fe). Note that C_P_ and ρ vary at cryogenic temperature, and adjustments were incorporated into SAR calculations. It was found that the SAR_Fe_ of EMG308 in M22 with and without LM5 at 35kA/m, 360kHz tested in 120kW RF coil is very similar (see Fig. S18E).

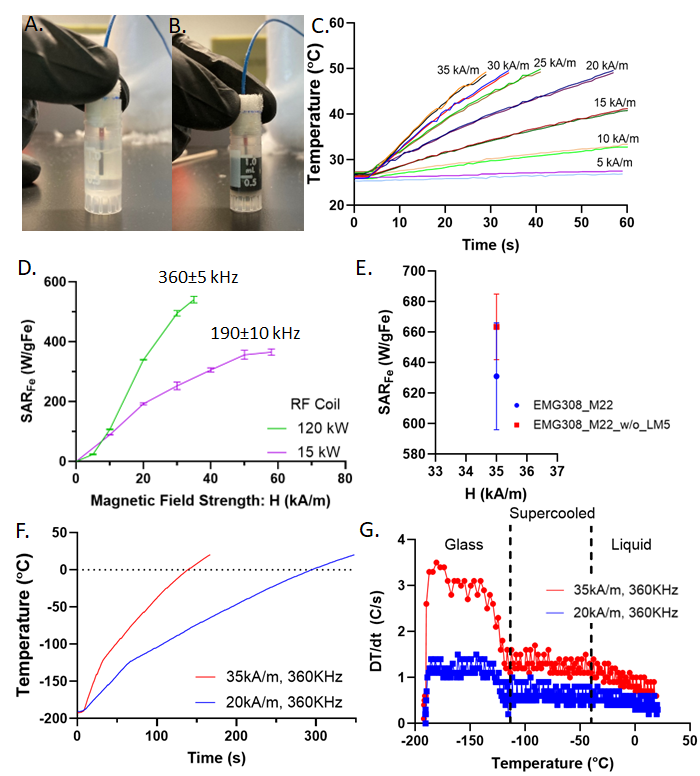

**Fig. S18: SAR measurement of CPA+IONPs**. Photos of A. CPA vial (control) with fiber optic temperature probe and B. CPA+IONP are in a similar setup to a SAR measurement control. C. Measured temperature vs. time curves used for SAR calculation of sIONP+M22 at 360kHz and varying magnetic field strength (listed next to each curve) in 120kW RF coil. D. SAR_Fe_ of EMG308 in water for varying magnetic field strength and frequency. E. SAR_Fe_ comparison for EMG308 in M22 with and without LM5 carrier solution. F. Temperature vs time curve and G. Slope (DT/dt) of temperature vs. time curve during cryogenic SAR measurement of sIONP in M22 at 35, 20 kA/m, and 360 kHz in 120 kW RF coil. The average of DT/dt in the three different temperature regions, i.e., glass, supercooled, and liquid, is taken to estimate cryogenic SAR_Fe_ as plotted in main Fig. 6B. Average C_p_ is used in these three distinct regions based on M22 literature values [39].

Eddy current heating estimation

Eddy current during nanowarming can be estimated using a simple correlation [36]:

$P_{eddy}=\sigma\left( \pi\mu_{o} \right)^{2}\left( r^{2} \right){f^{2}H}^{2}$ (11)

where H is magnetic field strength, f is frequency, r the radial distance from center of coil or the radius of sample, and µ_o_ is magnetic permeability of free space. It is to be mentioned that electrical conductivity σ in the above correlation should consider both dielectric loss (due to dipolar molecules) and DC electrical conductivity (due to ionic components) contributions.

$\sigma_{total}= \sigma_{dc}+\omega{\varepsilon^{''}\varepsilon}_{o}$(12)

In the past, dielectric properties and electrical conductivity were measured for different single CPA components. However, these values are limited and absent at lower RF frequencies, such as the kHz range where nanowarming coils operate (typically 100-500 kHz) for common organ vitrification CPAs and their mixture cocktails (only available for single component CPA such as glycerol at 1-100 kHz [40]). Now, in addition to dielectric heating due to polar molecules relaxation, at temperature above T_m_ of CPA cocktails, where mobility is increased due to less viscosity, another form of EM heating, i.e. ionic heating occurs, whose contribution to eddy heating is added via dc electrical conductivity of CPA solutions in above Eq. 15. This ionic contribution is due to the free ions present in the form of salts in the carrier solutions of CPA cocktails (LM5 for M22, Euro-Collins for VS55). This increases effective dielectric loss at near-zero temperatures and has been measured in literature at higher frequencies (~MHz) [41, 42]. Therefore, to capture the CPA cocktail behavior and in the absence of dielectric properties for organ vitrification CPAs (M22, VS55, 40%EG+0.6MSucrose), here, we assume dielectric properties from the closest available literature values on CPA+carrier solution (assumed values of σ & ε’’ as a function of temperature for 50%v/v DMSO + CPT at 27MHz as given elsewhere [41]). Fig. S19A shows the estimated value of total electrical conductivity of 50%DMSO from literature as ~0.1 S/m at >-20°C and decreases down to <0.02 S/m at ~-80°C [41] using Eq. 12 (See Fig. S19A). We calculated P_eddy_ using Eq. 11 for the 120 kW RF coil as a function of radius and magnetic field strength at 360kHz (Fig. S19B-C). It is found that P_eddy_ is ~25% (at >-20C) and reduces down to ~2% (at ~-80°C) of SAR_V_ i.e., % of nanoparticle heating (calculated assuming 10mgFe/mL IONPs at SAR_Fe_ ~ 1050 W/gFe in the cryogenic -90° to -40°C temperature range and SAR_Fe_ ~ 680 W/gFe in >-30°C temperature range) as shown in Fig. S19B.

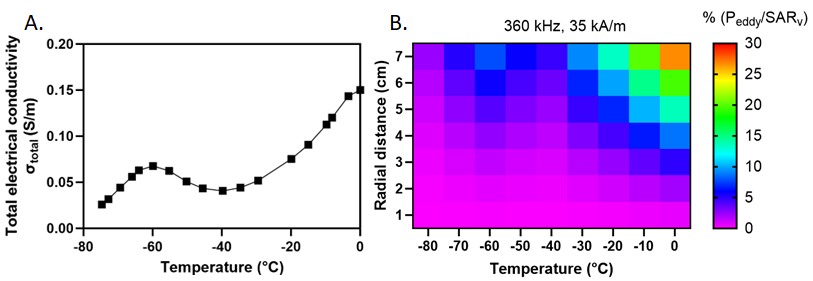

**Fig. S19: Eddy Current Heating Estimation.** A. Total electrical conductivity of a CPA variation at low temperature plotted using literature values from [41]. These conductivity values are used for P_eddy_ estimation (Eq. 15). B. Heat map plot of %P_eddy_/SAR_V_ with temperature and radial distance (from the center of RF coil) at a fixed magnetic field strength of 35 kA/m and 360 kHz. The red/orange region shows the highest value of P_eddy_, which occurs at larger radius and magnetic field strength near 0°C. The purple region shows that eddy heating is insignificant (<2% of SAR_V_) at lower radius (or field strength) at near zero temperatures and for all radii (or field strengths) at temperatures below <-40°C. SAR_V_ is calculated assuming 10mgFe/mL IONPs at SAR_Fe_ ~ 1050 W/gFe in the cryogenic -90° to -40°C temperature range and SAR_Fe_ ~ 680 W/gFe in >-40°C temperature range.

Nanowarming of multiscale CPA volumes

We also rewarmed 0.5L and 1L M22 volume at ~4.6 mgFe/mL EMG308, similar to 2L volume (Fig. S21A). As expected, the heating is similar for all three volumes, demonstrating that nanowarming is independent of size/volume (at a given IONP concentration). Warming rates achieved were ~85°C/min. We also rewarmed a 1mL cryovial containing VMP with EMG308 at ~100mgFe/mL after being vitrified in a LN2 bath. Rewarming was conducted from LN2 temperature of -196°C and stopped at near 0°C. The temperature was recorded every 0.2 sec for such a high concentration of IONP where ultra-fast warming rates were expected. Warming rates are plotted in Fig. S21 and calculated as ~1500°C/min in the 0 to -100°C range. Further, EMG308 in M22 at ~18mgFe/mL was also tested, showing warming rate dependence (approximately linear) on IONP concentration in CPA solutions (Fig. S21B). Note that the measurement response rate also becomes limited in fiber optics (data sampling time~0.5-1sec) when measuring rapid WRs (~1000°C/min).

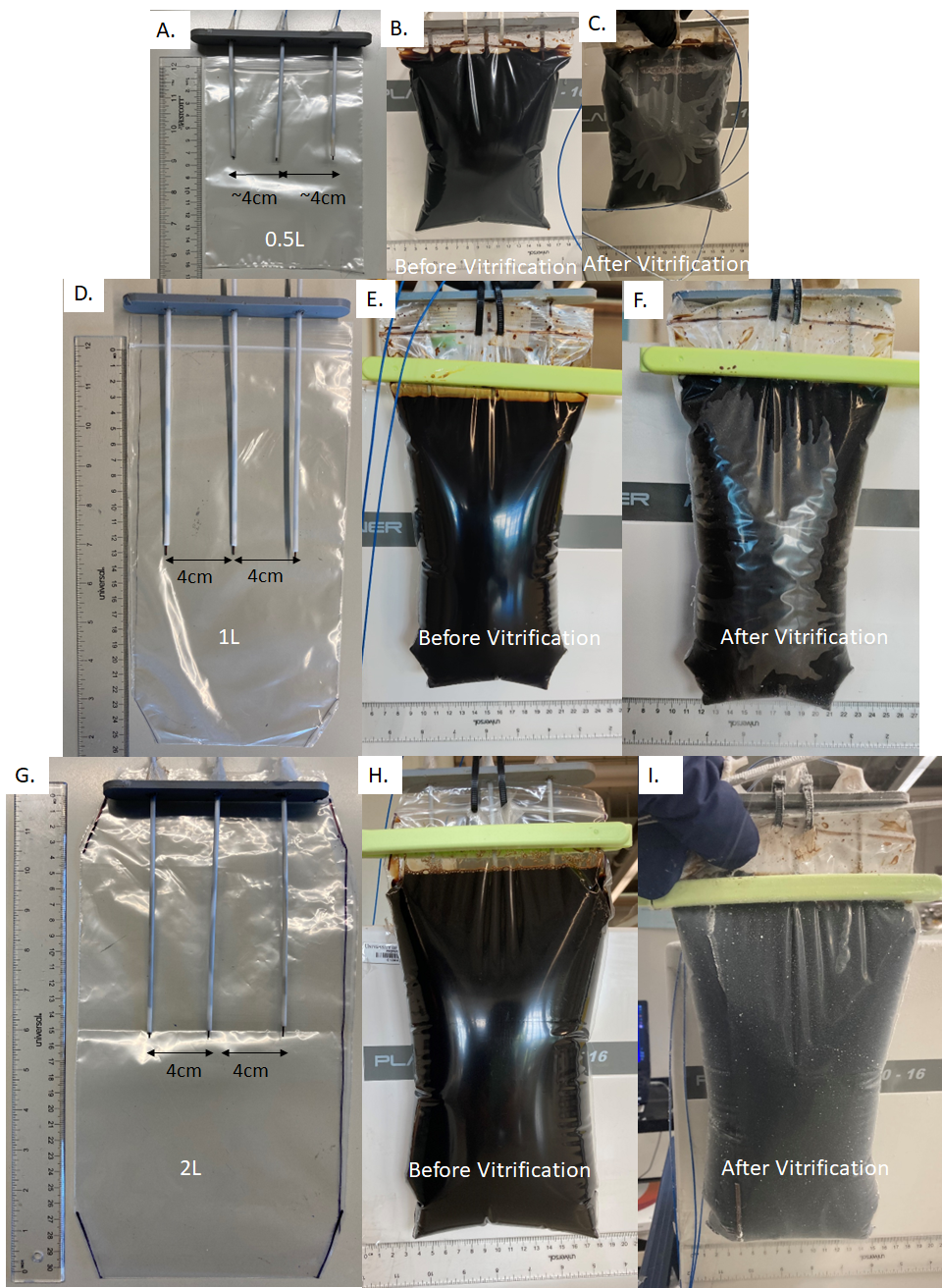

**Fig. S20: Photos of thermometry jigs for liter scale nanowarming**. EMG308+M22 cryobag with fiber optic probe placement for A. 0.5L, D. 1L volume and G. 2L volume. EMG308+M22 before and after vitrification B, C. 0.5L volume, E, F. 1L volume and H, I. 2L volume.

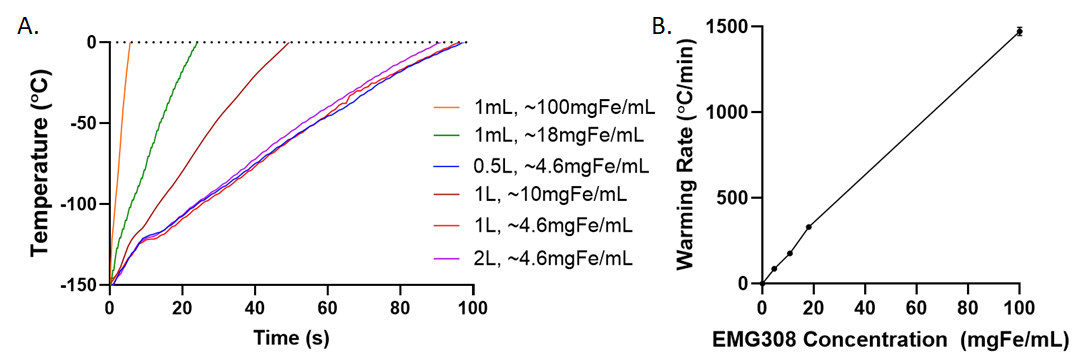

**Fig. S21: Nanowarming heating demonstration at multiple scales (mL to L)**. A. Thermometry (Temperature vs. time) for nanowarming of all the volumes tested: 1 mL at ~100mgFe/mL EMG308 in VMP, 1mL at ~18mgFe/mL EMG308 in M22 and 0.5L, 1L, 2L at ~4.6mgFe/mL EMG308 in M22 and 1L at ~10.4mgFe/mL EMG308 in M22 from the cryogenic vitrified state. B. Warming Rates as a function of IONP concentration. B. Average Warming rate (0 to -100°C) plotted as a function of IONP concentration for all the range of volumes rewarmed from the vitrified cryogenic state using nanowarming. The RF coil conditions were 35 kA/m and 360 kHz for all nanowarming volumes.

Characteristics length (L_C_), Cooling Rate, and Nanowarming Rate calculation for human organs

This section provides an approach to calculating the characteristic length of an organ, which can then be used to estimate cooling rates (assuming the organ alone with no surrounding volume in cryobags), as shown in Table S5. For example, we provide representative calculations of L_C_ for a human kidney whose Length ~ is 13cm, Width~ is 6cm, and Thickness~ is 3.5 cm, taken from the literature [43]. First, we assume an ellipsoidal kidney shape with semi-axes as half-length, half-width, and half-thickness.

$Volume of kidney (V)= \frac{4\pi}{3}*\frac{length}{2}*\frac{width}{2}*\frac{thickness}{2}$ (18)

$$Volume of kidney (V)= \frac{4\pi}{3}*\frac{13}{2}*\frac{6}{2}*\frac{3.5}{2}=142.9 {cm}^{3}$$

Using Knud Thomsen’s approximation for the surface area of an ellipsoid:

$Surface Area of kidney \left( A \right)\approx4\pi*\left[ \frac{\left( \frac{length}{2}*\frac{width}{2} \right)^{1.6}+\left( \frac{width}{2}*\frac{thickness}{2} \right)^{1.6}+\left( \frac{thickness}{2}*\frac{length}{2} \right)^{1.6}}{3} \right]^{\frac{1}{1.6}}$(19)

$$Surface Area of kidney \left( A \right)\approx4\pi*\left[ \frac{\left( \frac{13}{2}*\frac{6}{2} \right)^{1.6}+\left( \frac{6}{2}*\frac{3.5}{2} \right)^{1.6}+\left( \frac{3.5}{2}*\frac{13}{2} \right)^{1.6}}{3} \right]^{\frac{1}{1.6}}=161.9 {cm}^{3}$$

Now, the characteristic length for heat transfer becomes

$$L_{C}= \frac{V}{A}=\frac{141.9}{161.9}=1.08 cm$$

The center cooling rate can be calculated using a parametric fit of modeled CR vs L_C_ from the literature [24].

$\log\left( \frac{CR}{CCR} \right)\left[ \frac{^{\circ}C}{min} \right] =a+b*\log\left( L_{C} \left[ cm \right] \right)$ (20)

where a is 1.399 and b is -1.609 for M22, and the L_C_ of a human kidney is estimated to be 1.1 cm.

$$CR =0.4*[1.399-1.609*\log\left( 1.1 \right)]$$

$$CR =2.2 \frac{^{\circ}C}{min}$$

Lastly, we estimate the nanowarming rate (organ alone without any surrounding IONP volumes) in a human kidney with vascular fraction ~ 23% at 10mgFe/mL IONP concentration perfused in the vasculature as follows:

$\rho C_{P}\frac{\partial T}{\partial t}={SAR}_{V}$	(21)

plugging ${SAR}_{V} {=SAR}_{Fe}*C_{Fe}*Vascular fraction$

$Nanowarming rate=\frac{\partial T}{\partial t}=\frac{{SAR}_{Fe}*C_{Fe}*VF}{\rho C_{P}}$

$Nanowarming rate=\frac{1050 \left[ \frac{W}{gFe} \right]*10 \left[ \frac{mgFe}{mL} \right]*0.225}{1080 \left[ \frac{kg}{m3} \right]* 3300\left[ \frac{J}{kg}.K \right]}$

$Nanowarming rate \sim40 \frac{^{\circ}C}{min}$

Minimum vitrifiable CPA concentration calculation for a given characteristics length (L_C_)

First, characteristic length L_C_ for heat transfer is calculated for a geometry as discussed in above sections. Using the L_C_, the center (or minimum) cooling rate for a geometry can be estimated using the parametric fit (Eq. 20) of modeled CR vs L_C_ from the literature [24].

$\log\left( \frac{CR}{CCR} \right)\left[ \frac{^{\circ}C}{min} \right] =a+b*\log\left( L_{C} \left[ cm \right] \right)$ (20)

where CCR is 0.1°C/min, a is 1.399 and b is -1.609 for M22. Now using CCR vs. CPA concentration fitting from literature [44] as an approximation, for a CPA concentration one can estimate CCR or vice-versa as below.

$CCR \left[ \frac{^{\circ}C}{min} \right]={10}^{7}*e^{-0.269*Conc. [\%\frac{w}{w}]}$	(22)

Now, for minimum vitrifiable CPA concentration, achievable CR = CCR therefore rearranging Eq. 20 & Eq. 22 as follows.

$Min. Vitrifiable CPA Conc.[\%\frac{w}{w}]=(56.5+13.77*log L_{C}\left[ cm \right])$	(23)

Note that this is a first-order calculation and has inherent limitations such as CCR vs. CPA concentration fit was primarily developed for single component CPAs and therefore is limited in applicability for CPA cocktail mixtures such as M22 [44]. Furthermore, for simple approximation, we used M22 thermal properties as in the parametric fit of CR vs L_C_ and with varying CPA concentration thermal behavior (and hence CRs) would change to some extent [24].

**Table S1: Scale of systems successfully vitrified and rewarmed**.

**Scale**

**Timeline**

| **Volume** | **Biological system** | **Technique** | **Notable References (Author & Journal)** |
| --- | --- | --- | --- |
| 5 µL | Frog Spermatozoa | Vitrification and Convective Rewarming | [1, 2] |
| 0.2 mL | Mouse Embryos | Vitrification and Convective Rewarming | [3] |
| 0.2 mL | Human Ovarian Tissue | Vitrification and Convective Rewarming | [4] |
| 1-2 mL | Rabbit Jugular Vein & Articular Cartilage | Vitrification and Convective rewarming | [5, 6] |
| 5-10 mL | Sheep Ovaries | Vitrification and Convective rewarming | [7] |
| 10 mL | Rabbit Kidney | Vitrification and Convective Rewarming | [8] |
| 10-20 mL | Rat Heart + IONPs+CPA surrounding solution | Vitrification and Nanowarming | [9, 10] |
| 20-25 mL | Rat Kidney + IONPs + CPA surrounding solution |  | [11-13] |
| 25-30 mL | Rat Liver + IONPs + CPA surrounding solution |  | [14] |
| 50 mL | Rabbit Kidney | Vitrification and Dielectric Rewarming | [15] |
| 1-80mL | Porcine Arteries, Heart Valves, CPA+IONPs Solutions | Vitrification and Nanowarming | [16] |
| 80-90 mL | Porcine & Sheep Heart Valves + CPA surrounding solution | Vitrification and 2-stage Convective rewarming | [17, 18]  . |
| ~1L | Porcine Liver | Vitrification | “Current study” |
| 0.5-3L | CPAs (VS55, M22, EG+Sucrose) + IONPs | Vitrification and Nanowarming |  |
| 100mL -1.5L | Porcine &  Human Organs | Vitrification and Rewarming | To be demonstrated  (Future Cryopreservation Goal) |

**Table S2: CPA composition and calorimetric properties**.

| **CPA** | **CPA Concentration** | | |
| --- | --- | --- | --- |
|  | **VS55**  **(in EC)** | **M22**  **(in LM5)** | **40%v/vEG+0.6MSucrose**  **(in EC)** |
| **Total Concentration**  **(with carrier solution sugars*)** | 8.6 M  (55.3%w/w) | 9.5 M  (66.2%w/w) | 8.0 M  (60.4%w/w) |
| **Total Concentration**  **(without carrier solution sugars)** | 8.4 M  (52.6%w/w) | 9.3 M  (63.2%w/w) | 7.5 M  (57.9%w/w) |
| CCR, CWR (°C/min) | ~2.5, 50 [12] | 0.1, 0.4 [19, 20] | <1, ~45 [14] |
| T_m_: Melting Temperature (°C) | -38 [21], ~-45 [22] | ~-55 [23] | ~-43 [14] |
| T_g_: Glass Transition Temperature (°C) | ~-123 [21] | ~-123 [20, 22] | ~-121 [14] |
| **Components** |  |  |  |
| Dimethyl sulfoxide | 3.09 M | 2.86 M |  |
| Ethylene Glycol |  | 2.86 M | 7.15 M |
| Formamide | 3.08 M | 2.71 M |  |
| HEPES | 0.01 M |  |  |
| N-methyl formamide (NMF) |  | 0.51 M |  |
| Polyvinylpyrrolidone (2,500 kDa) |  | 0.01 M |  |
| Propylene Glycol | 2.21 M |  |  |
| Sucrose |  |  | 0.6 M |
| 3-Methoxy, 1,2-propanediol (MG) |  | 0.38 |  |
| X-1000 |  | 1% w/v |  |
| Z-1000 |  | 2% w/v |  |
| 5X EC carrier solution* | 20% v/v |  | 20% v/v |
| 5X LM5 carrier solution* |  | 20% v/v |  |

*Note: Carrier solution 1X Euro-Collins (EC) has ~0.2M Glucose, and 1X LM5 has 0.09M Glucose, 0.045M Mannitol, 0.045M Lactose. The carrier is prepared in a 5X concentrated solution to facilitate the required volumes. VS55 is ~62%w/v, M22 is ~71.6%w/v and 40%EG+0.6MSucrose is ~55.2%v/v.

**Table S3 Details of cryobag dimensions used in the liter-scale vitrification and nanowarming study.** All cryobags were purchased from McMaster Carr.

| Experiment Step | Volume | Height | Width | Material Thickness |
| --- | --- | --- | --- | --- |
| Cooling | 0.5 L | 6 inch (15.24 cm) | 6 inch (15.24 cm) | 0.051 mm |
|  | 1 L | 8 inch (20.32 cm) | 8 inch (20.32 cm) |  |
|  | 1 L  (Porcine Liver) | 10 inch (25.4 cm) | 8 inch (20.32 cm) |  |
|  | 3 L | 12 inch (30.48 cm) | 9 inch (22.86 cm) |  |
| Nanowarming | 0.5 L | 8 inch (20.32 cm)* | 5 inch  (12.7 cm) |  |
|  | 1 L | 10 inch (25.4 cm) | 8 inch (20.32 cm)* |  |
|  | 2 L | 10 inch (25.4 cm) | 8 inch (20.32 cm)* |  |

Note that these dimensions represent the cryobag without containing the sample (CPA, CPA+IONPs, or liver), and dimensions with the sample are listed in individual result figures. *Cryobags were heat-sealed to custom sizes (width in case of 1L, 2L nanowarming and height in case of 0.5L nanowarming) according to the volume tested and geometric constraints in control rate freezer (CRF) and 120 kW RF coil.

**Table S4: Thermo-physical properties of CPA used for computational FEM modeling.**

| CPA | Thermal Conductivity: k (W/mK) | | Specific Heat: Cp (KJ/kg.K) | Density: ρ (kg/m3) | Coefficient of Thermal Expansion: β *10^-4 (1/⁰C) | CCR (⁰C/min) | CWR (⁰C/min) |
| --- | --- | --- | --- | --- | --- | --- | --- |
| M22 | 0.3[25] | 3.43 (0⁰C)  3.378 (-18⁰C)  3.318 (-40⁰C)  3.180(-76⁰C)  3.324(-119⁰C)  1.461(-130⁰C)  1.318 (-149⁰C)  [26] | | 1100  [27] | 2.52 [28] | 0.1[19, 20, 29] | 0.4[19, 20, 29] |

**Table S5: Capabilities of custom-built state-of-art 120kW RF Coil System.**

| Magnetic Field Strength | 3 - 35 kA/m |
| --- | --- |
| Treatment Region Inner Diameter | 13.3 cm |
| Treatment Region Uniform Length | 20 cm |
| Field Uniformity in Treatment Region | +/- 5% |
| Frequency Range | 360 kHz +/- 10kHz |

**Table S6: Center cooling rates for human organs (no surrounding solution or container) based on characteristic lengths.**

| **Human Organs** | **Approximate Volume Adult Female-Male** | **Dimensions*** | **Lc**** | **Center/**  **Minimum Cooling Rate predicted***** |
| --- | --- | --- | --- | --- |
| Ovary | 4-6 mL [45] | 3, 2.5, 1.5 cm [46] | 0.35 cm | 14 °C/min |
| Eye | 6-7 mL [47] | 2.4, 2.34, 2.3 cm [47, 48] | 0.39 cm | 11.2 °C/min |
| Testis | 12-18mL [49, 50] | 4 ,3, 2.5 cm [51] | 0.50 cm | 7.6 °C/min |
| Pancreas | 60-80mL [52, 53] | 16, 4, 2 cm [54] | 0.54 cm | 6.7 °C/min |
| Kidney | 150-200 mL [55-58] | 13, 6, 3.5 cm [43] | 1.1 cm | 2.2 °C/min |
| Heart | 250-350 mL [56, 57] | 12, 8, 6 cm [59] | 1.3 cm | 1.6 °C/min |
| Lung | 400-600 mL [57, 58, 60] | 22, 8, 14 cm [61] | 1.9 cm | 0.9 °C/min |
| Brain | 1.3-1.5 L [56-58, 60] | 18, 13, 12 cm [62] | 2.2 cm | 0.7 °C/min |
| Liver | 1.5-1.8 L [56-58, 60] | 21, 15, 10.5 cm [63, 64] | 2.3 cm | 0.6 °C/min |

*Assuming adult human 64-70kg. There is a variation of ±10% between adult male vs. female and right vs. left paired organs based upon a literature review of these dimensions and volume.

**Assuming the organ is an ellipsoid with no surrounding volume in cryobag.

*** Using Eq. 20 from [24] where h of 100W/m^2^K was assumed for CRF.

Note: For lungs, the specified volume is tissue volume, whereas for L_C_ calculations, the volume used is total volume, including air space.

**Table S7: Nanowarming rates for organs loaded with 10mgFe/mL IONPs**.

| **Human Organs*** | **Approximate Volume Adult Female-Male** | **Lc** | **Estimated Vascular fraction** | **Predicted Nanowarming rate**  **@ 10mgFe/mL perfused conc.**** | **Conv. Warming Rate***** |
| --- | --- | --- | --- | --- | --- |
| Ovary [65-67] | 4-6 mL | 0.35 cm | 4-6% | 8.8°C/min | 15°C/min |
| Eye [68, 69] | 6-7 mL | 0.39 cm | 26-37% | 56°C/min | 12.5°C/min |
| Testis [70-72] | 12-18mL | 0.50 cm | 2-3% | 4.4°C/min | 8.5°C/min |
| Pancreas [73-75] | 60-80mL | 0.54 cm | 20-22% | 37°C/min | 7.5°C/min |
| Kidney [58, 76-78] | 150-200 mL | 1.1 cm | 20-25% | 40°C/min | 2.5°C/min |
| Heart [78-80] | 250-350 mL | 1.3 cm | 14-20%* | 30°C/min | 1.9°C/min |
| Lung [81, 82] | 400-600 mL | 1.9 cm | 34-36%* | 62°C/min | 1.0°C/min |
| Brain [58, 83] | 1.3-1.5 L | 2.2 cm | 3-8% | 10°C/min | 0.8°C/min |
| Liver [58, 78, 84] | 1.5-1.8 L | 2.3 cm | 13-28% | 36°C/min | 0.7°C/min |

*References in the organ column refer to vascular fraction calculations.

**IONP SAR_Fe_ is assumed to be 1050 W/gFe for calculations.

**Convective warming rate calculated for water bath rewarming at 37°C and h = 100 W/m^2^K based upon [10] and [24].

**Table S8: Nanowarming rates reported in cryopreserved volumes prior to this study.**

| Total Sample Volume | Nanowarmed sample | | ~Highest warming rate (WR) achieved | Temperature range WR is calculated | Reference |
| --- | --- | --- | --- | --- | --- |
| 0.5 mL | VS55+Co_35_Fe_65_ Nanowires | 10mg/mL | 1000°C/min | -150 to 0°C | [85] |
| 1 mL | VMP+EMG308 IONPs | 100mgFe/mL | 1500°C/min | -100 to 0°C | This study |
| 1 mL | HDF cells+VS55+EMG308/1200 | 59mgFe/mL | 200°C/min | -150 to -20°C | [86] |
| 20 mL | Mouse pancreatic islets+IONPs | 5mgFe/mL | 72°C/min | -123 to -38°C | [87] |
| 20 mL | VS55+sPION | 10mgFe/mL | 321°C/min | Not mentioned | [9] |
| 20 mL | Rat heart+VS55+sIONP | 10mgFe/mL | 75°C/min | Not mentioned | [10] |
| 25 mL | Rat kidney+VS55+sIONP | 10mgFe/mL | 65°C/min | -90 to -45°C | [12] |
| 35 mL | Rat liver+EG-Sucrose+sIONP | 10mgFe/mL | 61°C/min | -90 to -45°C | [14] |
| 40 mL | Porcine articular cartilage-OCA+CPAs | 2mgFe/mL | 77°C/min | -135 to -30°C | [88] |
| 80 mL | VS55+IONPs | 10mgFe/mL | 112°C/min | -135 to -30°C | [16] |
| 500 mL | M22+EMG308 | 4.6 mgFe/mL | 82°C/min | -100 to 0°C | This study |
| 1000 mL | M22+EMG308 | 10.7/4.6 mgFe/mL | 172/86 °C/min |  |  |
| 2000 mL | M22+EMG308 | 4.6 mgFe/mL | 88°C/min |  |  |
